# Tracking pre-mRNA maturation across subcellular compartments identifies developmental gene regulation through intron retention and nuclear anchoring

**DOI:** 10.1101/2020.10.23.352088

**Authors:** Kyu-Hyeon Yeom, Zhicheng Pan, Chia-Ho Lin, Hanyoung Lim, Wen Xiao, Yi Xing, Douglas L. Black

## Abstract

To globally assess the distribution and processing of gene transcripts across subcellular compartments, we developed extensive RNA-seq datasets of both polyA+ and total RNA from chromatin, nucleoplasm and cytoplasm of mouse ESC, neuronal progenitors, and neurons. We identified protein-coding genes whose polyadenylated transcripts were more abundant in chromatin than cytoplasm. We defined introns exhibiting cotranscriptional splicing, complete intron retention in cytoplasmic RNA, and many introns retained in nucleoplasmic and chromatin RNA but absent from cytoplasmic RNA, including new introns controlled during neuronal development. In particular, we found that polyadenylated *Gabbr1* transcripts are expressed in mESC but remain sequestered on chromatin until neuronal differentiation when they are processed and released to the cytoplasm. This developmental regulation of splicing and chromatin association demonstrates that the abundance of polyadenylated RNA is not always an indicator of functional gene expression. Our datasets provide a rich resource for analyzing many other aspects of mRNA maturation.

## INTRODUCTION

After transcription initiation, the nuclear maturation of pre-messenger RNA (pre-mRNA) requires splicing, polyadenylation, base modification, and release of the RNA from the chromatin template, before export from the soluble nucleoplasm to the cytoplasm for translation. For many genes, the bulk of expressed RNA exists in the cytoplasm as mature mRNA, while nascent, intron containing transcripts are limited to small nuclear puncta at the sites of transcription (Vargas et al., 2011; Coulon et al., 2014). For other genes, unspliced introns may remain after transcript completion but are ultimately excised to allow export (Girard et al., 2012; Popp and Maquat, 2013; Stewart, 2019). These nuclear transcripts are not necessarily found at their gene loci and their association with chromatin is not clear. Some polyadenylated Pol II transcripts, including many non-coding RNAs, are tightly associated with chromatin (Quinn and Chang, 2016). Although proteins have been identified that play roles in processes such as DNA template release, intron retention, RNA transport through the nuclear pore, and RNA targeting to nuclear decay pathways (Schmid and Jensen, 2018; Stewart, 2019), it is not well understood what features determine the chromatin, nucleoplasmic or cytoplasmic location of each transcript. The global distribution of RNA transcripts between subcellular compartments, their states of maturation, and how their maturation and localization change with development have not been well studied.

In earlier studies, we examined the relative kinetics of transcription, splicing, and nuclear export for macrophage transcripts induced by inflammatory stimuli (Bhatt et al., 2012; Pandya-Jones et al., 2013). This analysis used a subcellular fractionation procedure to enrich for nucleoplasmic or chromatin-associated RNA (Wuarin and Schibler, 1994; Pawlicki and Steitz, 2008; Pandya-Jones and Black, 2009; Khodor et al., 2012; Herzel and Neugebauer, 2015; Yeom and Damianov, 2017). The nucleoplasmic and chromatin compartments are operationally defined as the supernatant and pellet fractions, respectively, after nuclear lysis in a stringent buffer containing varying concentrations of NP-40, Urea, and NaCl. This solubilizes many components such as the U1 snRNP, while leaving other molecules almost entirely associated with the high molecular weight chromatin pellet (Wuarin and Schibler, 1994). The cytoplasmic fraction is enriched for mature mRNA, while the nucleoplasmic fraction contains recently matured transcripts that have been released from the chromatin template but have not yet reached the cytoplasm (Bhatt et al., 2012). This nuclear RNA pool can also include mature mRNAs associated with ER and mitochondria (Yeom and Damianov, 2017). The chromatin pellet is enriched for nascent RNA bound by elongating RNA Pol II, but this fraction also contains substantial polyadenylated RNA, including the Xist long non-coding RNA that is tightly bound to chromatin. This fraction also contains most of the Malat1 long non-coding RNA that is enriched in nuclear speckles that are adjacent to chromatin but only partially in close contact with it (Hutchinson et al., 2007; Fei et al., 2017). By following transcripts from inflammatory genes induced in macrophages, we found that partially spliced but polyadenylated transcripts in the chromatin fraction completed splicing over time, and were released to the soluble nucleoplasmic fraction before appearing in the cytoplasm as functional mRNAs (Bhatt et al., 2012). These studies focused on introns whose slow splicing impacted the rate of inflammatory gene expression. However, polyadenylated, partially spliced RNA has been long been observed in nuclei where its interactions and localization are largely unknown.

The consequences of intron retention are diverse and complex to dissect. A number of studies have shown that splice sites and binding of spliceosomal components can prevent nuclear RNA export (Hautbergue, 2017; Stewart, 2019; Garland and Jensen, 2020). Nevertheless, some intron containing transcripts are exported to the cytoplasm as alternative mRNA isoforms that either encode an alternative protein or be subject to altered translation and decay (Jacob and Smith, 2017; Wegener and Müller-McNicoll, 2018). Other introns that are slow to be excised relative to transcription can be ultimately removed and their transcripts exported as fully spliced mRNAs (Ninomiya et al., 2011; Bhatt et al., 2012; Hao and Baltimore, 2013; Pandya-Jones et al., 2013; Frankiw et al., 2019a). The slower kinetics of these introns can determine the rate of overall gene expression and can be modulated in response to cellular stimuli. Such transcripts can create a nuclear pool of partially spliced RNA, which acts as a reservoir to feed the cytoplasmic mRNA pool upon splicing activation. A group of these introns found in genes affecting growth control and cell division have been called “detained introns” to distinguish them from classical “retained introns” found in cytoplasmic mRNA (Boutz et al., 2015; Braun et al., 2017). A similar pool of incompletely spliced transcripts affecting synaptic function is found in neurons, where cell stimulation induces their processing and export to allow changes in mRNA pools independent of transcriptional kinetics (Mauger et al., 2016). The term “retained intron” thus encompasses a wide range of molecular behaviors and regulatory mechanisms.

Retained introns are also more difficult to characterize than other patterns of alternative splicing in whole transcriptome RNA-seq data, where overlapping patterns of alternative processing can be mis-called as intron retention by sequence analysis tools (Wang and Rio, 2018; Broseus and Ritchie, 2020). Many RNA-seq studies have identified conditions leading to higher levels of unspliced introns across the transcriptome (Wong et al., 2013; Braunschweig et al., 2014; Schmitz et al., 2017; Jacob and Smith, 2017; Parra et al., 2018). These studies have not always distinguished between nuclear and cytoplasmic RNA or examined the ultimate fate of the partially spliced transcripts, information that is essential to understanding the biological role of such regulation and its mechanisms.

Here we undertook a broad examination of how RNAs are distributed between subcellular compartments and how this compartmentalization changes with development. We distinguished transcripts in the nucleoplasmic and chromatin-associated RNA pools from cytoplasmic mRNAs to identify RNAs exhibiting different localization behaviors and assessed their state of processing. Focusing on fully transcribed polyadenylated RNA, we defined groups of introns enriched in different subcellular compartments. Some transcripts containing retained introns are exported to the cytoplasm as mRNAs. Other introns are present at high levels in the nucleus but have been excised from the cytoplasmic RNA. Finally, the polyadenylated products of some genes are abundant as partially spliced transcripts in the nucleoplasmic and particularly the chromatin fractions, but are absent from the cytoplasm, spliced or not. The RNA localization and splicing of these gene transcripts are regulated to allow differential gene expression.

## RESULTS

### Both coding and non-coding RNAs exhibit defined partitioning between cellular compartments

To broadly categorize RNAs enriched in different cellular compartments, and gain insight into how this compartmentalization might be regulated across cell types, we generated deep RNA-seq data from mouse embryonic stem cells (mESC), a neuronal progenitor cell line derived from embryonic mouse brain (mNPC), and explanted mouse cortical neurons cultured *in vitro* for 5 days (mCtx) (Figure 1A). RNA was isolated from three fractions of each cell: cytoplasm, soluble nucleoplasm, and chromatin pellet using the procedures previously described (Wuarin and Schibler, 1994; Pandya-Jones and Black, 2009; Bhatt et al., 2012; Yeom and Damianov, 2017). The quality of subcellular fractionation was assessed by immunoblot for GAPDH and TUBULIN proteins as cytoplasmic markers, U1-70K protein fractionating with the soluble nucleoplasm, and HISTONE H3 as a chromatin marker (Supplemental Figure S1A).

**Figure 1.**
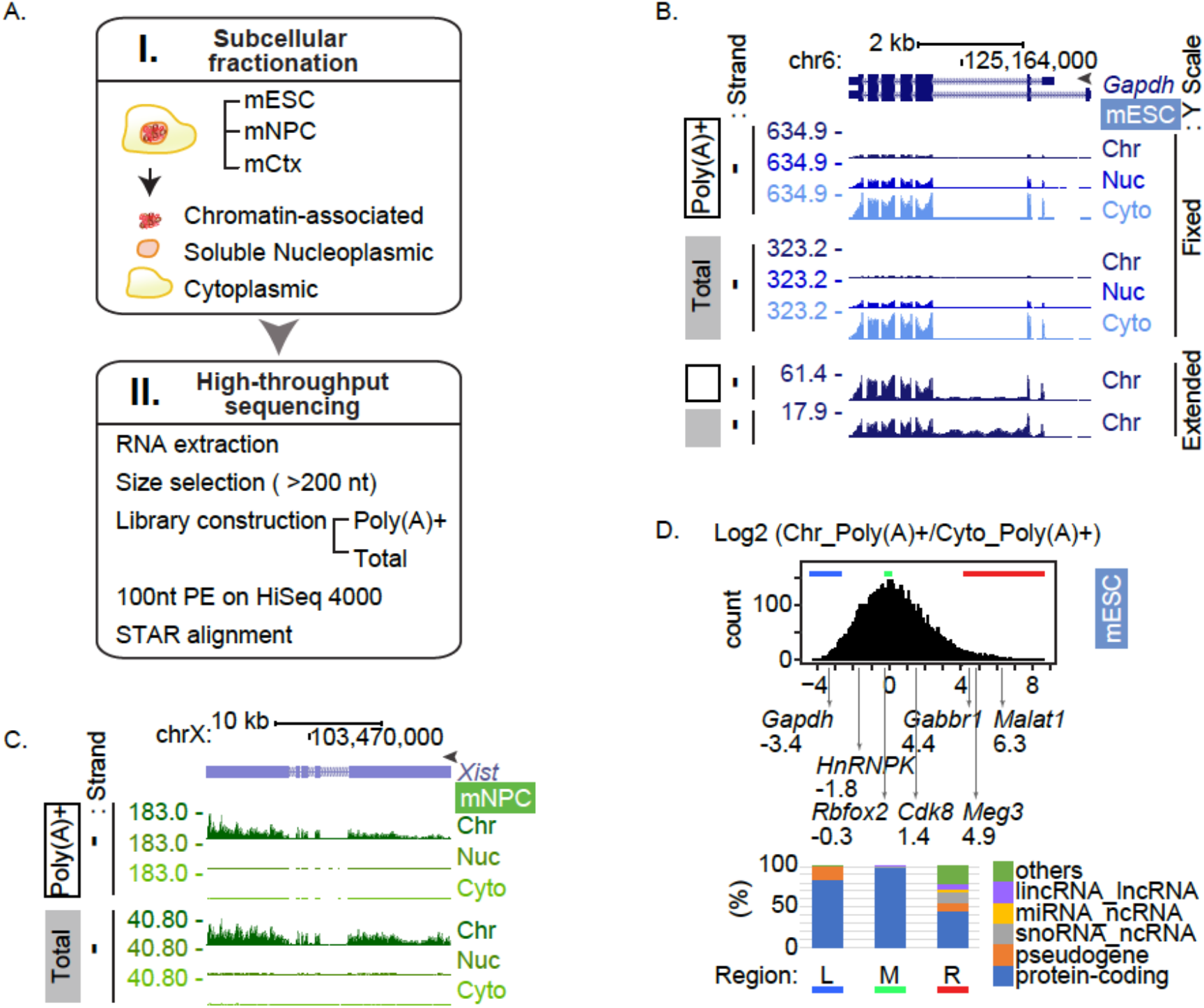
RNA partitioning between subcellular compartments. A. Workflow used in this study. I. Mouse ESC, NPC, and cortical neurons (mCtx, 5DIV) were fractionated to separate cytoplasm, soluble nucleoplasm, and chromatin bound material. II. Both total (ribominus) and polyA+ RNA was isolated from each fraction and sequenced to generated 100 nt paired end reads, which were aligned to the genome B. Genome browser tracks of the *Gapdh* locus in mESC. Isoforms from ENCODE/GENCODE gene annotation version M11 are diagrammed at the top. Poly(A)+ RNA (open box) and Total RNA (grey box), as well as the peak RPM are noted on the left. RNA from the chromatin-associated (Chr), soluble nucleoplasmic (Nuc), and cytoplasmic (Cyto) fractions are labeled at the right. The fixed Y scale (RPM) shows the strong enrichment of Gapdh RNA in the cytoplasm. The bottom two tracks show chromatin RNA with an extended Y scale to observe the intron reads. C. Genome browser tracks of the *Xist/Tsix* locus in the female mNPC line. Xist expression of female mNPC line. The fixed Y scale shows strong enrichment of Xist RNA in the chromatin fraction. D. Distribution of chromatin partition indices. The Chromatin / Cytoplasm ratio of the averaged read counts of each gene was determined and transformed to log2 using DEseq2 (log2 [Chr_Poly(A)+ / Cyto_Poly(A)+]). Genes expressed above the median TPM of Chr_Poly(A)+ RNA plotted as a distribution along the log2 scale. The partition index of representative genes are indicated under the plot. The biotypes of the 400 genes from bottom (left (L), blue bar), peak (middle (M), green bar), and top (right (R), red bar) of the distribution are presented in the bar graph below. The “other” biotype category includes additional ncRNAs and RNAs not yet defined for biotype.

The three fractions from three cell types yielded RNA from nine conditions. To provide information on the maturation of transcripts in each condition, RNA was isolated from each fraction as two separate pools. A total RNA pool depleted of ribosomal RNA (Total) will include nascent incomplete transcripts. In contrast, a polyadenylated pool (Poly(A)+) includes RNAs whose transcription and 3’ processing is complete. Each RNA pool from each fraction was isolated from three separate cultures of each cell type to yield biological triplicates of each experimental condition. These RNA pools were converted to cDNA libraries, sequenced on the Illumina platform to yield 100 nt paired end reads, and aligned to the genome (Supplemental Table S2). Gene expression markers for each of the three cell types confirmed the expected patterns of ESC, NPC or neurons (Supplemental Figure S1B). Clustering of gene expression values across all the datasets showed the expected segregation by cell type, fraction, and replicate, for both the poly(A)+ and total RNA libraries (Supplemental Figure S1C). The resulting 54 datasets constitute an extensive resource for examining multiple aspects of RNA maturation and its modulation during development [GEO:in process]. In addition to the libraries we describe in this study, we also generated libraries of small RNAs from all samples. These were partially described in an earlier paper and can be used to assess miRNA maturation and other processes (Yeom et al., 2018). These 9 datasets are also available from [GEO:in process].

Examining read distributions in the different libraries and fractions, we found that the housekeeping gene *Gapdh* (Figure 1B) yields similar patterns of reads from either the poly(A)+ or the total RNA populations, with the RNA being most abundant in the cytoplasm. The total Gapdh RNA on chromatin contains intron reads from the nascent transcripts (Figure 1B bottom). Although more abundant in the soluble nucleoplasm and especially in the cytoplasm, polyadenylated Gapdh transcripts are also found in the chromatin fraction, but in contrast to the total RNA these lack intron reads. We also examined the long non-coding RNA Xist, which condenses on the inactive X chromosome in female cells (Figure 1C). The mNPCs were isolated from female mice, and Xist is seen to partition almost completely to chromatin in these cells. The poly(A)+ and the total RNA samples yielded very similar patterns of Xist reads indicating that this RNA is largely spliced and polyadenylated (Brockdorff et al., 1992). Other non-coding RNAs yielded more complex patterns of subcellular partitioning that changed with cell type. The paraspeckle lncRNA Neat1 is more highly expressed in mESC than mNPC or neurons (Supplemental Figure S2A). The short polyadenylated form (Neat1_1) predominates in ESC and is found mostly with chromatin but also in the nucleoplasm. The longer non-polyadenylated Neat1 RNA (Neat1_2) is seen in the total RNA samples, and is also chromatin-enriched. Whether this is a stable long isoform or nascent RNA is not clear. Interestingly this longer RNA contributes a larger portion of the Neat1 transcripts in mNPC and neurons, consistent with observations that Neat1 cleavage and polyadenylation may be modulated in cells to control the production of the short isoform (Naganuma et al., 2012). Overall, we find that gene transcripts can exhibit diverse patterns of enrichment and processing across the different fractions and cell types.

Because the relative transcript numbers and overall library complexity will differ between fractions, RPM (reads per million) values or other read number normalizations of individual genes cannot be directly compared between different subcellular fractions. Using qRT-PCR in mESC to directly quantify individual transcripts in different fractions, we found that for cytoplasmic enriched transcripts in both the poly(A)+ and the total RNA libraries, RPM values undercounted the RNA abundance in the cytoplasmic fraction relative to the chromatin and nucleoplasm (Supplemental Table S3). On the other hand, for RNAs that are primarily chromatin associated, qRT-PCR quantification yielded cytoplasmic to chromatin ratios that were similar to relative RPM numbers (Supplemental Table S3). Although the absolute transcript levels were not quantifiable by RPM, the ratios of these RPM values did reflect their relative enrichment in each fraction across a variety of genes. As an index for how RNAs partitioned between the chromatin and cytoplasmic pools, we used DESeq2 (Anders and Huber, 2010) to measure the fold change in reads for each gene between the chromatin and cytoplasmic poly(A)+ RNA. This returns the ratio of the averaged read counts for each gene between fractions. For genes whose TPM value in chromatin was over the median and which had read counts greater than 0 in the cytoplasm (13,036 genes), this chromatin partition index was distributed over a 100 fold range centered on 1 (Log_2_=0). Thus, a typical gene showed equal read counts in chromatin and cytoplasm (Figure 1D). Examining the Ensembl annotations (V.91) for genes in the left, middle, and right side of this distribution (400 genes each), we found that genes with predominately cytoplasmic reads as well as genes with roughly equal read numbers in cytoplasm and chromatin were annotated almost entirely as protein-coding genes. For example, on the left edge (Figure 1D), Gapdh RNAs partition much more strongly to the cytoplasm than is typical. In the middle of the distribution, Rbfox2 RNAs exhibit slightly fewer reads on chromatin than in the cytoplasm, whereas Cdk8 exhibits 2 to 3 fold more chromatin reads (Figure 1D). Thus, although the transcripts from protein coding genes are usually most abundant in the cytoplasm, a substantial fraction of a gene’s RNA product is often found in the nucleus and is chromatin-associated. Comparing qRT-PCR quantification for select genes to their chromatin partition indices, we found that RNAs from genes exhibiting a partition index above 3.6 were actually more abundant in chromatin than the cytoplasm. This included about 3% of protein coding genes. At the right edge of the curve, the 400 genes producing the mostly highly chromatin enriched transcripts included the expected non-coding RNAs, such as primary-miRNAs, snoRNAs, and lncRNAs. Interestingly, this group also included a substantial number of protein-coding genes, including *Clcn2* and *Ankrd16* (shown in Supplemental Figures S2C and S2D), and *Gabbr1*, which is analyzed further below. For these protein-coding genes, the majority of the polyadenylated product RNA is chromatin associated where it is presumably inactive for protein expression (Supplemental Figures S2C and S2D).

Examination of individual genes whose poly(A)+ transcripts remain sequestered with chromatin showed that their splicing was modulated across cell types. The chromatin-associated Meg3 non-coding RNA is well expressed in mESC and neurons but not in mNPC (Supplemental Figure S2B). Meg3 is the host transcript for the microRNAs miR-770 and miR-1906-1. Mature miR-770, processed from the last Meg3 intron, is neuronally expressed (data not shown). Interestingly this intron is absent from the RNA in mESC where it is apparently more efficiently spliced. By contrast in neurons, this intron is abundant in the chromatin fraction of polyadenylated RNA, where its reduced excision might allow more efficient processing of miR-770 (Supplemental Figure S2B). This is consistent with observations that Drosha cleavage occurs on chromatin and that perturbations that cause a host transcript to be released from chromatin reduce miRNA expression (Pawlicki and Steitz, 2008; Liu et al., 2016a). We previously used some of this mESC data to examine expression of primary miR-124a-1 in mESC and how its Drosha processing is blocked by PTBP1 in the chromatin fraction (Yeom et al., 2018). For Meg3, the processing of miR-770 by Drosha may be modulated instead by the excision rate of its host intron. The upstream portion of Meg3 that includes miR-1906-1 undergoes complex processing events and exhibits more splicing in neurons than in mESC. This may indicate that an additional product from the gene, possibly miR-1906-1, is also differentially regulated between ESC and neurons. These introns are present in the polyadenylated RNA and are not more abundant in the total RNA than the surrounding exon sequences, indicating that there is not an excess of excised intron, which could also give rise to the miRNAs. Overall the data indicate that splicing of the Meg3 transcript is regulated on chromatin to allow differential expression of its mature products.

### Chromatin associated transcripts can be spliced either co-transcriptionally or post-transcriptionally

It is expected that most introns will be transient species within the chromatin RNA, with many introns excised prior to transcript completion, while some introns with slow kinetics will be removed later. Various studies estimate that 45 to 84 percent of introns are cotranscriptionally excised in mammals (Ameur et al., 2011; Bhatt et al., 2012; Girard et al., 2012; Khodor et al., 2012; Tilgner et al., 2012; Windhager et al., 2012). Several approaches compare read numbers for spliced (exon-exon) and unspliced (exon-intron or intron-exon) junctions in nascent RNA to those in total RNA to measure cotranscriptional excision (Tilgner et al., 2012; Windhager et al., 2012; Herzel and Neugebauer, 2015). To ensure that measurements are of the nascent RNA, this requires removal of polyadenylated RNA from the chromatin fraction and prevents parallel analysis of posttranscriptional events. Other studies noted the presence of sawtooth patterns in RNA-seq data from total cellular RNA, where reads peak in exons and then steadily decline to the next exon or recursive splice site. Such a pattern is thought to indicate that the time needed to excise an intron is small relative to the time for RNA synthesis through the next intron downstream (Ameur et al., 2011; Duff et al., 2015; Sibley et al., 2015). We also observed sawtooth read densities on certain introns in the total chromatin RNA pools (data not shown). As reported previously, these sawtooth read patterns were infrequent and are lost on introns shorter than 50kb (Ameur et al., 2011) (data not shown). Since the large majority of introns removed prior to transcript termination do not present such a sawtooth pattern, it would be useful to establish other tools for simultaneously defining both co- and post-transcriptional introns from the same libraries.

As an alternative for defining cotranscriptional and posttranscriptional intron excision, we compared the total RNA from chromatin to the poly(A)+ RNA from the same fraction. Introns remaining in polyadenylated RNA must be excised after transcription or be dead-end products. For example, in the *Sorbs1* gene (Figure 2A) reads are observed across all the introns in the total RNA from chromatin indicating the presence of unspliced introns in the nascent transcripts. In the polyadenylated RNA on chromatin, reads are largely absent from introns indicating that by the time of polyadenylation or shortly after, these introns have been spliced out. However, one intron in Sorbs1 exhibits substantial read numbers in poly(A)+ RNA on chromatin that are reduced in RNA from the nucleoplasm and absent from the cytoplasm (Figure 2A). This intron is presumably excised after cleavage/polyadenylation. While most introns are absent from the polyadenylated RNA and likely spliced cotranscriptionally, there are many transcripts with one or more introns that are highly retained in the polyadenylated chromatin associated RNA (Figure 2A and 2B). The comparison of intron levels in total and poly(A)+ RNA on chromatin provides a simple bioinformatic metric for distinguishing co-versus post-transcriptional excision.

**Figure 2.**
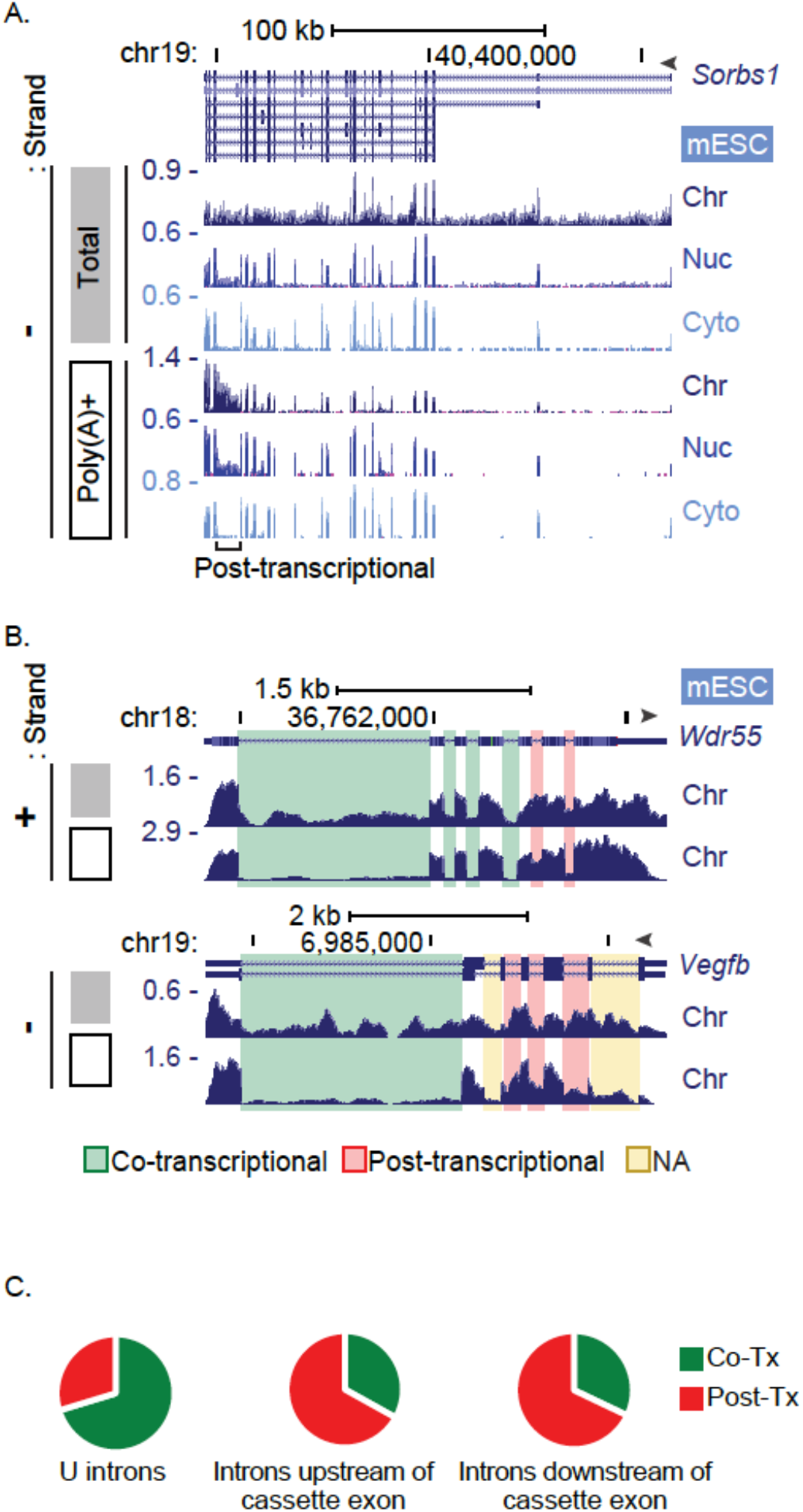
Co-transcriptional and post-transcriptional intron excision. A. Genome browser tracks of the *Sorbs1* locus in mESC. The total RNA (grey box) in the chromatin fraction shows reads in all the introns, but the polyA RNA (open box) is limited to exon sequences except for the third to last intron. This intron is defined as posttranscriptional using the criteria described in the text. B. Genome browser tracks of chromatin-associated RNA mapping to the *Wdr55* and *Vegfb* loci in mESC. Total (grey box) and poly(A)+ (open box) are shown. Cotranscriptionally and posttranscriptionally spliced introns are highlighted in green and red respectively. Yellow highlighted introns were not analyzable due to multiple processing patterns. C. Proportions of co- and posttranscriptional splicing events in mESC. A total 49,692 U introns passed filters for expression, lack of 3’ bias, and sufficient junction reads to determine FI. These were identified as cotranscriptionally (Co-Tx, green) or posttranscriptionally (Post-Tx, red) excised by comparing FI values in the total and poly(A)+ RNA on chromatin, using the criteria described in Supplemental Figures S3C – S3E. Introns upstream (2,779) and downstream (2,744) from simple cassette exons were analyzed in the same way.

To compare intron levels in the total and poly(A)+ RNA pools, we determined fractional inclusion values (FI; Fraction of intron Inclusion, Supplemental Figure S3A) by counting reads across exon-intron, intron-exon, and exon-exon junctions. Assessing intron retention (IR) by FI value can be confounded by alternative splicing, polyadenylation or transcription initiation events occurring within the intron being measured (Supplemental Figure S3B). To avoid errors in IR measurements arising from other processes, we defined a set of introns exhibiting a unique Ensembl v91 annotation without alternative processing events (Supplemental Figure 3B). This set of 149,333 “unique” introns (U introns) across 28,733 genes was used for subsequent analysis. Focusing on the mESC RNA, we determined the FI values of all U introns in the total RNA and the poly(A)+ RNA for genes above the median expression level as measured by Kallisto (Bray et al., 2016). We included only introns excised by the major spliceosome with GU/AG splice junctions. Reads from poly(A)+ RNA can be biased toward the 3’ ends of transcripts containing long unspliced introns. To avoid undercounting in the poly(A)+ samples, we removed genes where reads per nucleotide length from the second exon were less than half that of the second to last exon. To filter out introns that were not measurable due to anomalies in the generation of particular junction reads, we removed introns yielding a FI value below 0.1 in the total RNA, and introns with a zero value for one or more of the junction read counts. In mESC, these criteria returned 49,629 U introns within 7,672 genes for analysis.

Of the 49,629 U introns being measured, 34,939 introns (within 6,952 genes) exhibited low FI values in the poly(A)+ RNA (FI < 0.1), and are presumably spliced before transcript completion. Conversely, 14,753 introns within 5,550 genes exhibited a FI value greater than or equal to 0.1 in the poly(A)+ RNA. These introns (29.7 %) appear to be excised posttranscriptionally, with many highly unspliced in the chromatin poly(A)+ RNA despite being fully spliced in other fractions. By this analysis, at least 70.3 % of introns within our analysis set are excised cotranscriptionally, similar to estimates made by other methods (Figure 2C, Supplemental Figures S3C – S3F, and Supplemental Table S4). On the other hand, the majority of genes (5,550 out of 7,672) have at least one post-transcriptionally spliced intron. Restricting the analysis to the top quartile of expressed genes rather than the top half, the fractions of co and posttranscriptional splicing change only slightly (70.7 % co-transcriptional). The fraction of cotranscriptionally spliced introns is also essentially the same if the analysis is restricted to the first introns in each transcript or to internal introns. For introns that are the last intron transcribed before the polyA site, a slightly higher fraction is classified as posttranscriptional, presumably because they are polyadenylated more rapidly after intron synthesis (Supplemental Figure S3F). Thus, posttranscriptional splicing does not appear to be associated with higher or lower gene expression, or with the position of an intron along the gene. Examples of introns defined as co or posttranscriptional by these measures are shown in Figure 2B. Although in the minority, posttranscriptionally spliced introns are found across a wide range of genes, and often exhibit high FI values in the chromatin fraction, even though the cytoplasmic RNA is completely spliced.

In addition to the U introns analyzed above, we also analyzed a set of introns flanking simple cassette exons that could also be unambiguously measured for FI. Using the same parameters to define co-versus posttranscriptional splicing, we found a reversal in the percentages. Of these introns flanking alternative exons, approximately 67 % exhibit high read numbers (FI>0.1) in the poly(A)+ RNA and thus appear to be excised posttranscriptionally (Figure 2C and Supplemental Figure S3E). This was seen for introns both upstream and downstream of the cassette exon. These data indicate that the majority of regulated splicing events occur with slower kinetics than the excision of typical constitutive introns.

### Retained introns can be classified by their enrichment in the chromatin, nucleoplasmic, and cytoplasmic compartments

A variety of fates are possible for transcripts that retain introns after polyadenylation. Intron containing transcripts can be sequestered in the nucleus until they are spliced, or can undergo nuclear decay. Other intron containing mRNAs are exported unspliced to the cytoplasm where they can be translated or undergo NMD. To categorize introns based on both their retention levels and location, FI values for the unique intron set in the polyadenylated RNA of all cells and fractions were subjected to X-means cluster analysis (Figure 3A and https://xinglabtrackhub.research.chop.edu/intron_retention/) (Pelleg and Moore, 2000). Interestingly in all three cell types, the clustering algorithm defined four groups of introns. The largest cluster Group A, containing 49,981 introns in mESC, was almost entirely spliced in all three fractions. Introns in Group B (7,529), exhibited measurable retention in the poly(A)+ RNA from chromatin, but showed nearly complete splicing in the nucleoplasm and cytoplasm (Figure 3A). Group C introns (1,351), including introns in Zfp598 and Neil3 (Figure 3B), showed higher FI values in the chromatin and nucleoplasm than Group B, but were almost completely excised from the cytoplasmic RNA. The smallest cluster of only 247 introns in mESC, Group D, was almost entirely retained in all three fractions. Each of the other two cell types also generated four clusters with similar splicing levels and similar numbers of introns in each group (Figure 3A).

**Figure 3.**
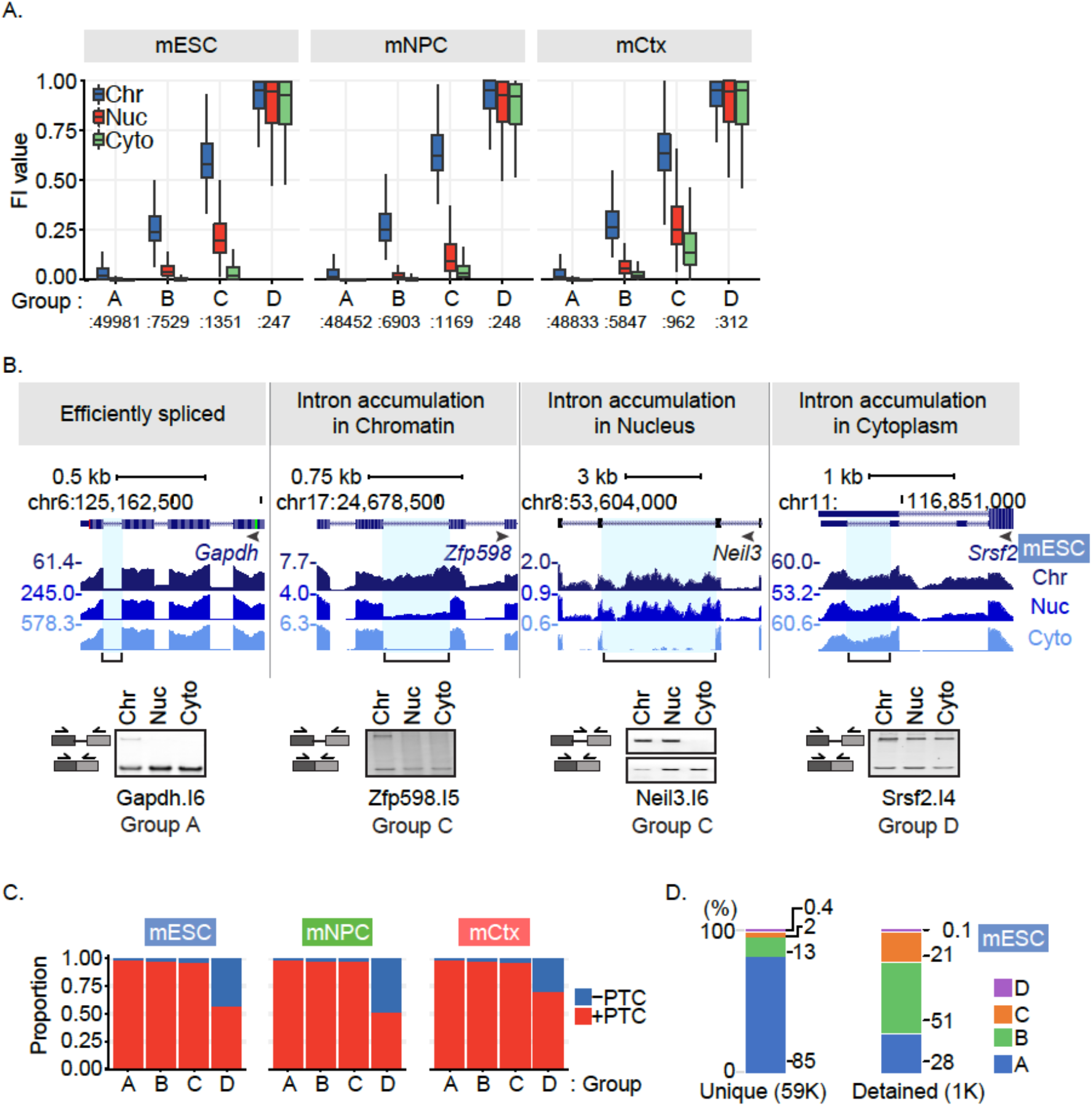
Intron Groups defined by their retention level and fractionation behavior. A. X-means clustering was applied to intron FI values and fraction enrichment in mESC, mNPC, and mCtx. The FI distribution of introns in each subcellular fraction for each group are shown. Group A introns are efficiently spliced intron in all fractions, while Group D introns unspliced in all fractions including the cytoplasm. Groups B and C show intermediate levels of splicing in the nucleus and chromatin but are fully spliced in the cytoplasm. B. Genome browser tracks (top) and RT-PCR validation (bottom) of representative transcripts in mESC. Validated introns are indicated by a blue highlight and a bracket below. Representative gel images are from 3 biological replicates. C. The proportion of introns containing a premature termination codon (PTC) in frame with the upstream sequence is shown for each cluster in each cell type. D. Proportion of introns in defined groups. Clustering analysis was performed on 59,108 uniquely annotated introns (U introns) from mESC and assigned to Groups A, B, C, or D. The U intron set from mESC included 1,000 previously defined as detained introns, which are enriched for introns in Groups B and C. The percent of each group in the intron set is shown.

Group B and C introns that do not leave the nucleus can be seen to have different properties from Group D introns that also have high retention levels in the cytoplasm. A much larger percentage of Group D introns are found in 5’ and 3’ UTR sequences, where they will not disrupt the primary reading frame, but will likely affect translation and decay (Supplemental Table S5B). Group D introns were also found to be depleted of in-frame premature termination codons (PTC) compared to Groups A, B and C (Figure 3C), presumably due to selection to prevent NMD in the cytoplasm. These observations indicate that the different intron clusters arise from selection for different functions and roles for the intron containing RNAs.

We found that in transcripts where all the introns were in the U category and could be assigned a group (Supplemental Figure S3G), those RNAs containing at least one Group C intron have a higher average chromatin partition index than transcripts with no Group C intron (Supplemental Figure S3H). Previous work defined a set of nuclear enriched transcripts in mESC containing what are called detained introns (DI), whose splicing is modulated in cancer and growth control pathways (Boutz et al., 2015; Braun et al., 2017). Of 3,150 detained introns, 1,021 were on our U intron list (Supplemental Tables S5A and S5B). 21 introns did not pass expression filter, leaving 1000 introns to be compared to the intron groups A, B, C, D, and these predominantly fall into Groups B and C, in agreement with the earlier studies (Figure 3D). However, these 1,021 detained introns were only a subset of the nearly 9,000 retained introns identified in Groups B and C (Supplemental Table S6B). Similar to the detained introns and the delayed inflammatory gene introns identified previously, these new retained introns likely affect additional cellular functions by maintaining a nuclear pool of unspliced RNA and altering the movement of material through the gene expression pathway.

### Predicting retained introns

To examine whether introns in different groups could be identified by their sequence features alone, we developed a deep learning model for predicting intron behavior. We extracted 1,387 sequence features from the first and last 300 nucleotides of each intron and from two flanking exons (Supplemental Table S7A). For introns less than 300 nucleotides, the intron interval includes some adjacent exon sequence. Analyzed features included short motif frequencies, predicted RBP binding elements, propensity to form local secondary structure, splice site strength scores, conservation scores, and nucleosome positioning scores. This feature information was used to train a three-layer deep neural network tasked with predicting whether an intron belonged in Group A, B, C, or D (Figure 4A).

**Figure 4.**
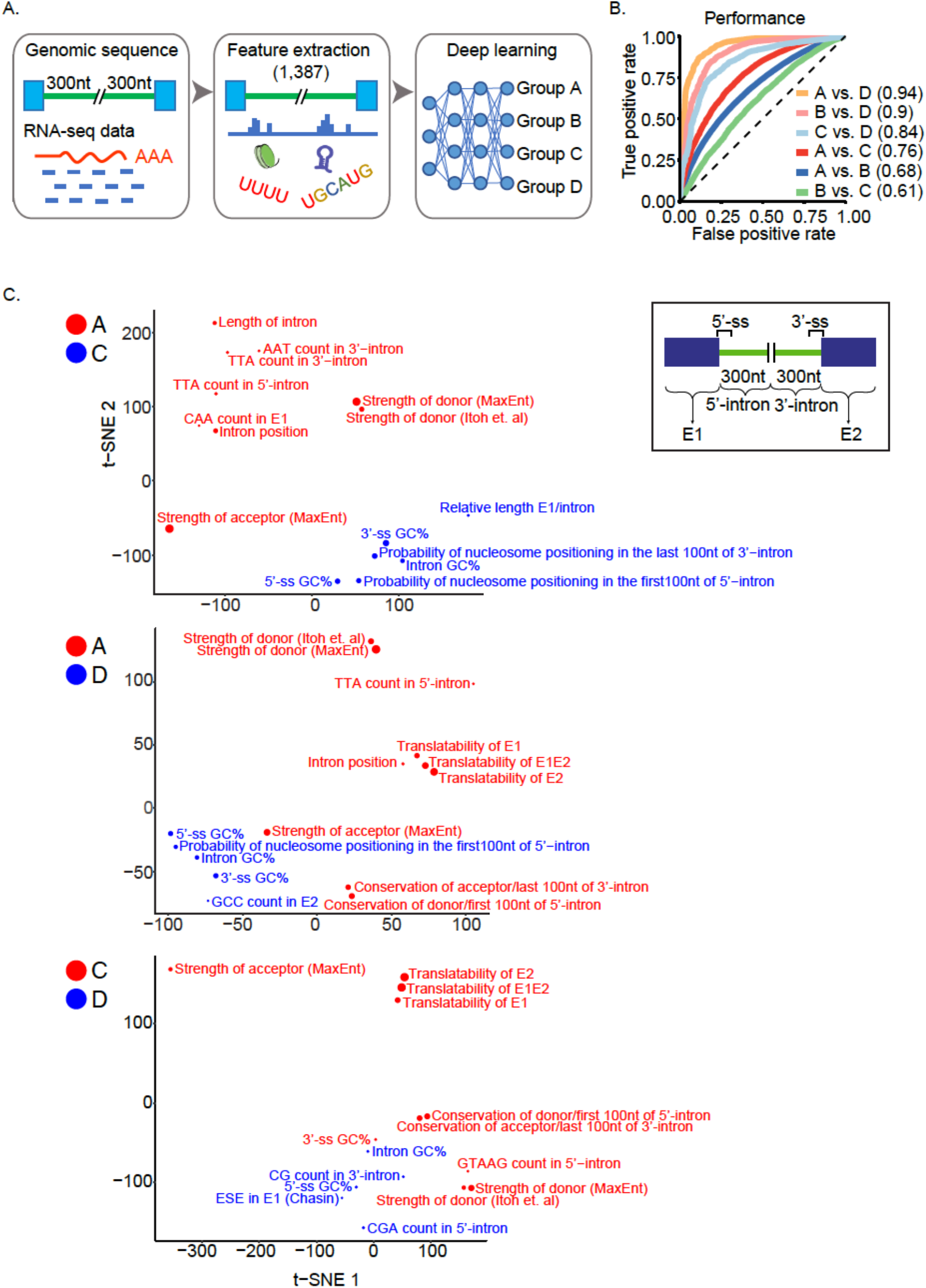
Deep Learning Analysis of Intron Groups. A. Flow diagram for training the deep neural network. B. Performance of the model in distinguishing introns of different groups. Receiver Operating Characteristic curves were plotted for individual pairwise comparisons. Area under the curve (AUC) values are shown in parentheses. C. The 15 genomic features most predictive for distinguishing intron groups. A t-distributed stochastic neighbor embedding algorithm was used to plot these features onto 2 dimensions. The strengths of the regulatory features were defined by the decrease of performance on held-out data when the values of each regulatory feature were substituted by their median. Features distinguishing Group A from Groups C and D are shown above and those distinguishing Group C from Group D are below. Features are colored blue or red according to the group for which they are positively correlated.

The performance of the model was assessed using Receiver Operating Characteristic (ROC) curves plotting the false and true positive rates (Figure 4B). The model was highly predictive in distinguishing Group D introns from A, yielding an Area Under the Curve of 0.94 (AUC = probability that any true positive will rank higher than any true negative). Group D introns could also be distinguished from Group B and C (AUC=0.9 and 0.84, respectively), and Group B and C introns from Group A with reduced accuracy (AUC = 0.68 and 0.76, respectively). Thus, the Group D introns are most different from the introns of other groups.

To assess the features of Group C and D introns that distinguish them from each other and from Group A, we isolated the top 15 features predictive of intron retention or its absence and used a t-distributed stochastic neighbor embedding algorithm (t-SNE) to project them onto two dimensions (Figure 4C and Supplemental Table S7B for top 50 features). As previously observed, high splice site strength scores were predictive of Groups A and C over D, and also Group A over C. Other features redundant with splice site strength scores were also predictive of Groups A or C, including GTAAG count in the 5’ portion of the intron and the conservation of the splice site sequences. Features predictive of Group D over A included the predicted propensity for nucleosome positioning in the intron. Translatability of the flanking exons and their spliced product was predictive of Groups A and C over D. This may reflect a greater percentage of Group D introns in 5’ and 3’ UTR sequences (Supplemental Table S5B). Conversely, the translatability of the exon-intron-exon unit containing the retained intron was predictive of Group D over Group C, in agreement with the Group D introns being depleted of in frame termination codons (Figure 3C) and adding a coding segment to the mRNA. Overall, the data indicate that intron retention is controlled by many factors each having relatively small effect.

We examined whether particular sequence elements correlated with the intron group assignments, indicative of regulatory protein binding sites. The model did not clearly identify known elements affecting nuclear localization or intron retention such as constitutive transport elements or decoy exons (Li et al., 2006; Parra et al., 2018). However, the sequence conservation score of the 5’ portion of the intron was predictive of Group D over Groups C or A, and conservation of both ends of the intron was predictive of C over A. Particular triplet motif frequencies within introns or their flanking exons were also predictive of intron behavior. For example, CGA triplets in the 3’ portion of the intron were predictive of Group D over C, whereas TTG and GTT triplets in the 5’ intron segment were predictive of Group C over D. The predictive power of intron sequence conservation and of multiple triplets indicate that particular RNA/protein interactions likely determine the retention properties of these groups.

### Intron retention and chromatin association are regulated with neuronal development

In mESC, mNPC and neurons, the X-means analysis yielded four clusters of introns based on their level of splicing in the different compartments. These cluster definitions allow bioinformatic analysis of IR regulation between cell types. While many introns maintain their classification between cell types (Figure 5A, left side), some introns switched their group (Figure 5A, right side). One example of this was Med22 (Figure 5B), which contains a highly retained intron 3 (I3) in all three fractions of mESC (Group D). This intron became more spliced in mNPC and was classified as Group C, and then became almost fully spliced as a Group B intron in neurons, while intron 1 (I1) was maintained as a Group A intron in all three cell types. Med22 encodes a protein subunit of the transcriptional mediator complex. The retained Med22 intron is the last intron in the transcript and its retention or splicing creates MED22 proteins with different C-terminal peptides. It will be interesting to examine how the virtually complete switching between these isoforms affects mediator function in the different cell types.

**Figure 5.**
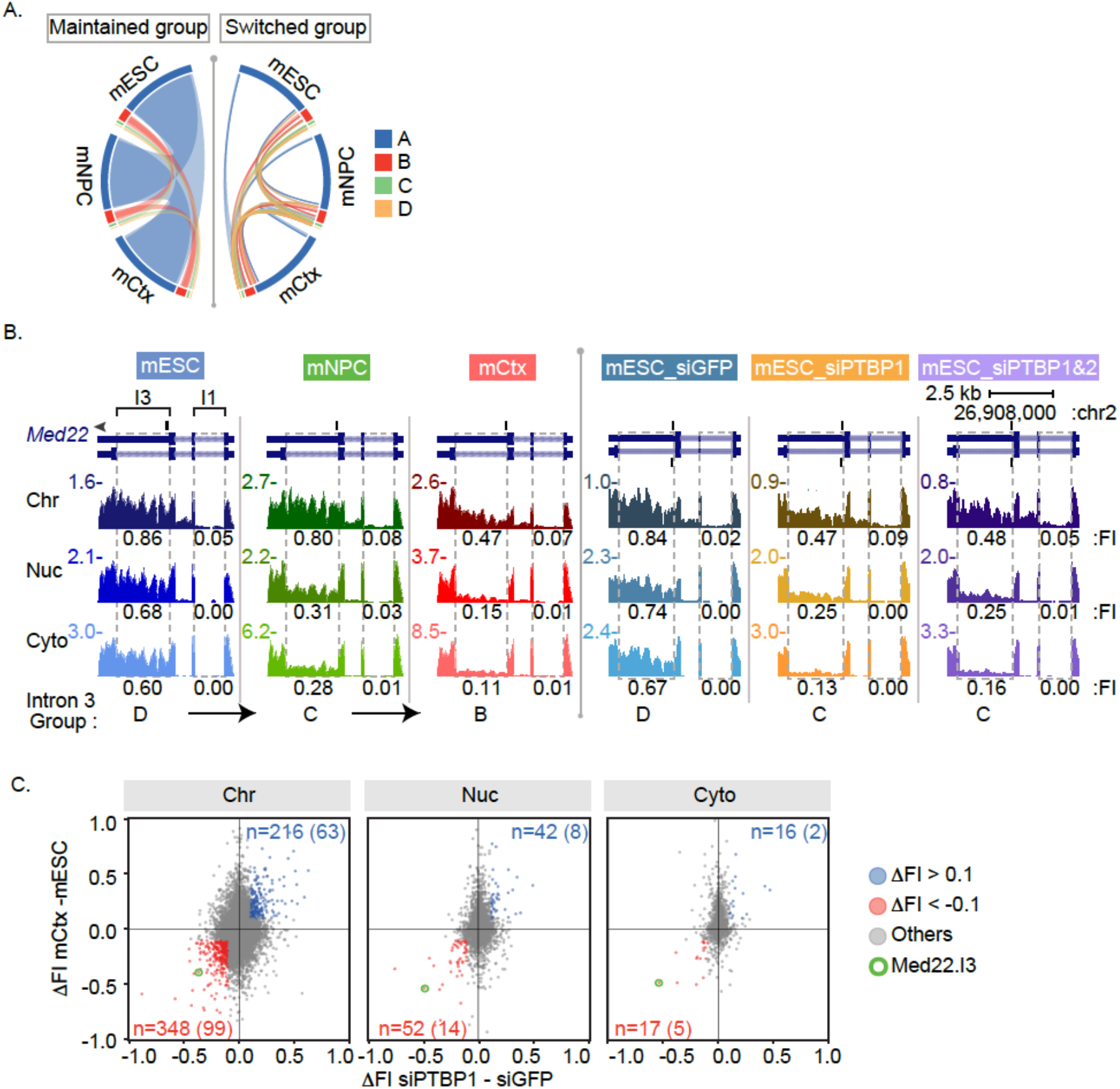
Regulation of intron retention and chromatin association during neuronal development. A. Circos plot of intron group changes between cell types (mESC, mNPC, and Cortical Neurons, mCtx). Introns that did not change groups are displayed on the left side. Introns that switched group between cell types shown on the right side. B. Genome browser tracks of *Med22* during neuronal differentiation (left three panels) and after PTBP knockdown (KD) in mESC (right three panels). Dashed boxes indicate introns in our U intron set with measured FI values (introns 1 and 3). FI values are shown under each track. Group classification of intron 3 is shown at the bottom. Intron 1 was classified as Group A in all cases. C. Scatter plots for each fraction of FI change between mESC and neurons (mCtx) plotted against FI change after PTBP KD in mESC. Introns with delta FI less than −0.1 in both conditions are marked in red, and introns with delta FI greater than 0.1 in blue. Number of introns showing these changes are shown for each fraction with the bracketed number reporting the number with PTBP1 iCLIP tags in mESC (Linares et al., 2015). Intron 3 of Med22 is circles in green.

As a set, the group switching introns are presumably part of the extended alternative splicing programs that are modulated during neuronal development. Examining their functions in Gene Ontology analyses, we found that 231 genes containing highly spliced introns in mESC that became unspliced in neurons (switching from Group A or B to Group C or D) were enriched in processes such as ribosome biogenesis, organelle assembly, and metabolism. These functional categories may reflect the different proliferation rates and metabolic status of the two cells. In contrast, 413 genes whose introns were unspliced in mESC and became more spliced in neurons (switching from Group C or D to Group A or B) were enriched in GO biological processes of glutamatergic synaptic transmission and organelle localization by membrane tethering, in keeping with the gene expression and cell morphological changes that accompany the transition to the early neuronal state (Supplemental Figure S4).

The changes in splicing between mESC, mNPC and neurons are driven by changes in the expression of multiple protein regulators. In previous work, we and others have characterized extensive alternative splicing programs controlled by the polypyrimidine tract binding proteins, PTBP1 and PTBP2 (Keppetipola et al., 2012; Vuong et al., 2016). In ESC and other cells, PTBP1 maintains alternative splicing patterns characteristic of non-neuronal cells, and PTBP1 downregulation is a key step in neuronal differentiation. Our earlier work on splicing programs regulated by PTBP1 in ESC focused on cassette exon changes in genes affecting neuronal development (Linares et al., 2015). However, PTBP1 regulation of retained introns, including the Med22 intron, have been described in a neuronal cell line (Yap et al., 2012). We wanted to examine whether additional targets of PTBP1 could be identified in ESC by characterizing the RNA of the chromatin compartment.

To assess the extent of PTBP1 regulation on chromatin associated RNA, we fractionated cells after PTBP1 knockdown and measured the splicing of polyadenylated RNA in the different compartments by RNA-seq. This confirmed the PTBP1 dependent regulation of Med22 intron 3, causing it to shift from a Group D intron to Group C (Figure 5B, Right Side). Focusing on retained introns, we find that many more splicing changes are identified in the chromatin associated RNA than in the nucleoplasmic and cytoplasmic fractions (Figure 5C). As shown previously with cassette exons, the PTBP1 dependent introns were seen to change with neuronal differentiation as PTBP1 levels drop (Figure 5C). These include introns identified previously and many more (Yap et al., 2012) (data not shown). Other introns whose splicing changes with neuronal development but are not sensitive to PTBP1 are presumably regulated by other factors.

By examining the chromatin associated RNA, our analysis identified substantially more PTBP1 regulated and other retained introns than previously recognized. The transcripts containing these introns may remain in the nucleus similar to detained introns, or may be exported to the cytoplasm and then lost to NMD. To assess this, we used data from a previous study of unfractionated polyadenylated RNA after UPF1 knockdown that globally identified NMD targets in mESC (Hurt et al., 2013). A majority of Group A, B and C introns are predicted to induce NMD, if their parent transcripts were exported to the cytoplasm (Figure 3C). However, we find that of 871 genes containing PTBP1 dependent retained introns identified in the chromatin fraction, only 87 of these genes exhibit greater than 10% transcript upregulation after Upf1 depletion and are likely NMD targets (Supplemental Table S6C). Thus, the majority of the PTBP1 dependent retained intron transcripts likely stay in the nucleus and will be eliminated by nuclear RNA decay pathways.

Looking more broadly at whether NMD might create the apparent nuclear enrichment of some transcripts, we found that protein-coding genes with high chromatin partition indices were actually less likely to show increases after Upf1 depletion than other genes across the distribution (Supplemental Table S6D). For the genes in the L, M, and R regions in Figure 1D, NMD targets constituted 4.2, 7.2, and 1.1 percent respectively. We suggest that rather than NMD causing the observed nuclear enrichment by depleting the cytoplasmic RNA, the nuclear enrichment may buffer the effect of NMD on the level of total RNA. It would be interesting to examine the effect of Upf1 knockdown specifically on the levels of cytoplasmic mRNA.

### Developmental control of gene expression through intron retention and nuclear sequestration

We previously identified genes for neuronal proteins, including *Ptbp2* and *Dlg4*, that are expressed in ESC’s but generate an NMD sensitive isoform preventing expression of protein (Zheng et al., 2012). These genes are broadly transcribed but post-transcriptional regulation maintains their developmental specific protein expression. Here we identify new transcripts enriched in the chromatin fraction of mESC’s (Figure 1D). While some of these are seen to be NMD targets, many are only mildly or not affected by Upf1 depletion (Supplemental Table S6D). For these RNAs, there are apparently additional processes that prevent their expression as mRNAs. A notable example is Gabbr1, which encodes the GABA B receptor 1, an inhibitory neurotransmitter receptor whose cytoplasmic mRNAs are highly expressed in neurons, more moderately expressed in mNPC, and found at only low levels in the mESC (Figure 6A). By immunoblot, GABBR1 protein is only observed in neurons (Figure 6C). Strikingly, in the chromatin fractions of mESC the *Gabbr1* precursor RNA is present at high levels that nearly match those seen in mNPC and neurons (Figure 6A). Although this RNA is polyadenylated and most introns have been excised, it shows only limited splicing of several introns, and the RNA is not fully mature. One region encompassing introns 4 and 5 exhibits a complex pattern of alternative processing in neurons but is largely unprocessed in mESC (Figure 6A). Gabbr1 RNA increases ~1.8 fold with Upf1 depletion from mESC. In addition to this NMD effect, Gabbr1 mRNA expression is apparently also inhibited by a combined process of splicing inhibition and sequestration on chromatin. Upon differentiation into neurons, the chromatin partition index of Gabbr1 RNA shifts from 4.43 to −0.69, as the RNA becomes fully processed and is released from chromatin to allow its appearance in the cytoplasm as mature mRNA (Figure 6A).

**Figure 6.**
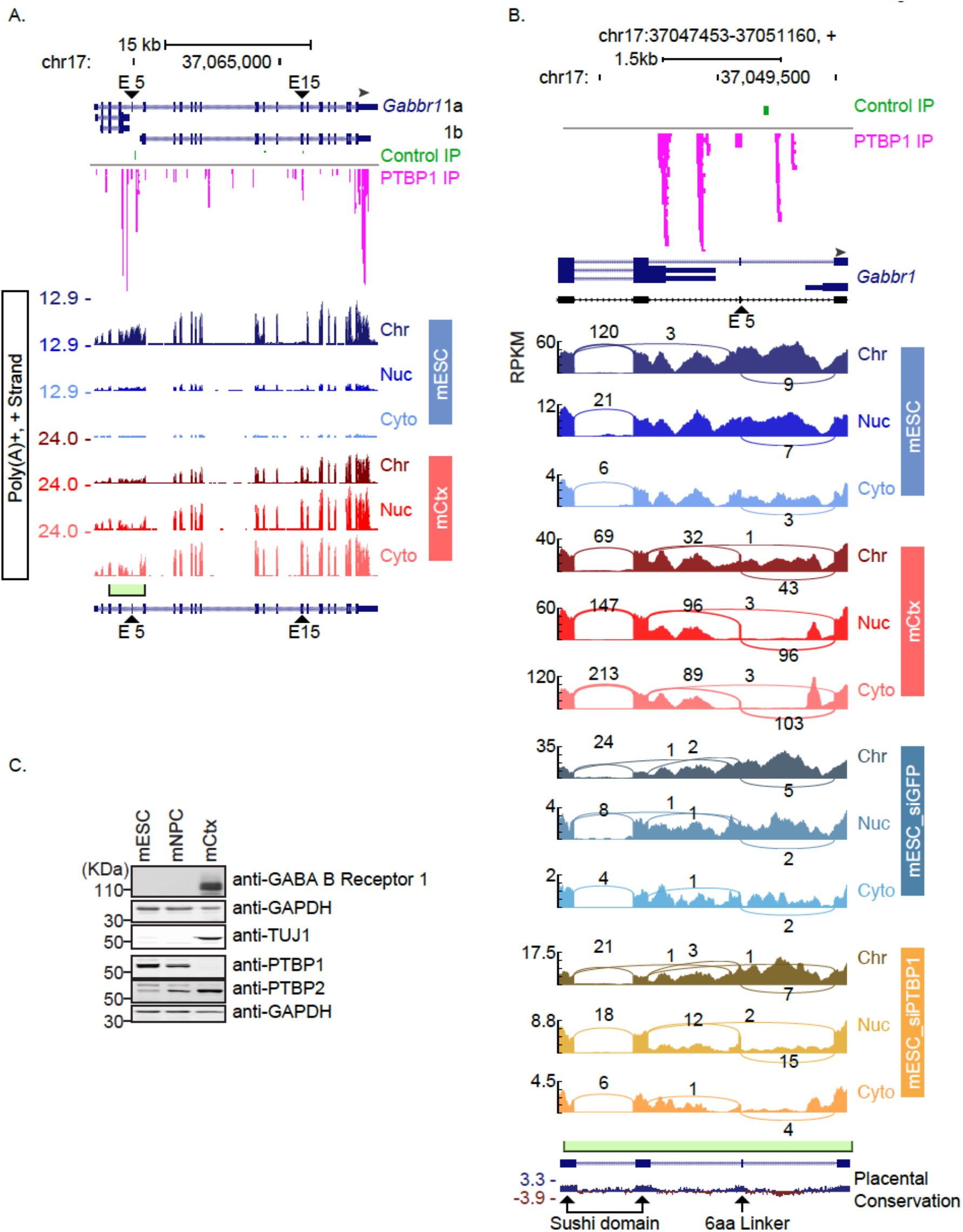
Chromatin enrichment and PTBP1 regulation of Gabbr1 transcripts. A. Genome browser tracks of *Gabbr1* in mESC and mCtx. PTBP1 iCLIP tags in mESC are plotted above the RNA-seq tracks. The Y axis indicates the maximum RPM in each cell type. The green box and bracket marks the intron 4-5 region expanded in panel B. PTBP1 responsive exons 5 and 15 are marked with arrowheads. B. Sashimi plots of the Gabbr1 intron 4-5 region in mESC, mCtx, and after PTBP knockdown in mESC. RPKM is plotted on the Y axis. PTBP1 responsive exon 5 is marked with an arrowhead. Exons encoding the two sushi domains and the 6aa linker are marked on the conservation track at the bottom. C. Immunoblot showing expression of GABBR1 protein relative to other proteins in mESC and cortical neurons.

Previous work found that PTBP1 regulates exon 15 in the Gabbr1 transcript in a neuronal cell line (Makeyev et al., 2007). To assess the intron 4-5 region, we examined iCLIP data mapping PTBP1 binding sites in mESC (Linares et al., 2015). This confirmed PTBP1 binding upstream of *Gabbr1* exon 15 and to the *Gabbr1* 3’ UTR. Besides the 3’ UTR, the most prominent PTBP1 binding was seen in several peaks in the intron 4-5 region (Figure 6A and 6B). Examining the fractionated RNA-seq data after PTBP1 siRNA treatment, we found that PTBP1 depletion led to processing of the Gabbr1 RNA into the neuronal isoforms, including activation of exon 15 and activation of the exon 5 microexon encoding a 6 amino acid linker of Gabbr1a (Figure 6A and 6B). Interestingly, although a greater fraction of the now processed Gabbr1 RNA was present in the cytoplasm after PTBP1 depletion, there was more of this spliced RNA in the soluble nuclear fraction, and a majority of the RNA was in the chromatin fraction and still unprocessed in the intron 4-5 region. This occurred despite the fact that exon 15 was strongly activated for splicing by the loss of PTBP1 (Makeyev et al., 2007), even in the chromatin fraction (Supplemental Figure S5B). The partition index was 2.25 after PTBP1 depletion. GABBR1 protein was not seen to be increased in mESC after PTBP1 depletion (data not shown). Thus, although PTBP1 strongly affected the processing of Gabbr1, its depletion did not yield the predominantly cytoplasmic distribution of RNA seen in neurons or its translation. There are likely additional factors mediating the chromatin sequestration of the RNA in mESC. Although highly transcribed in ESC, *Gabbr1* expression is blocked by a combination of splicing repression, NMD of the transcripts that enter the cytoplasm, and sequestration of the unprocessed RNA on chromatin, with the latter mechanism having the largest effect.

## DISCUSSION

### A Resource for the Analysis of RNA-level Gene Regulation

We developed extensive sets of RNA sequencing data to examine RNA maturation events across cellular location and developmental state. Applying these data to analyze intron retention, we compare total and polyadenylated RNA across subcellular fractions and cell types to define classes of introns exhibiting different regulatory behaviors, and we uncover a novel form of gene regulation acting on chromatin associated RNA. We find that a substantial fraction of the polyadenylated RNA product of some genes is incompletely spliced and still associated with chromatin. This points to a limitation for whole transcriptome measurements of gene expression that assess total cellular polyadenylated RNA; The RNA being measured in these studies is not all cytoplasmic mRNA. The presence of nuclear polyadenylated RNA may thus contribute to the observed lack of correlation between RNA and protein levels in global gene expression measurements (Edfors et al., 2016; Liu et al., 2016b).

The isolation of chromatin associated RNA has frequently been used to enrich for nascent pre-mRNAs and other short lived species (Pandya-Jones and Black, 2009; Davidson et al., 2012; Herzel et al., 2017). We find that many introns are only observed in the total RNA of this fraction, while others are also present in the polyadenylated RNA. Quantifying this difference, we estimate that 70% of introns within our analysis set are spliced before the RNA has been completely transcribed. Although this roughly agrees with other studies, we believe it is a lower-bound estimate in our system because the criteria for counting cotranscriptionally excised introns required a measurable presence of the intron in the total RNA. Interestingly, we find that introns flanking alternatively spliced cassette exons are mostly spliced posttranscriptionally - exhibiting significant intron retention levels in the polyadenylated RNA. These introns may be spliced more slowly than typical constitutive introns because of the complex regulatory RNP structures that must assemble onto the sequences flanking alternative exons. By creating a pool of unspliced RNA for these genes, the delayed splicing may allow additional controls over the isoform choice. It will be interesting to examine whether the subset of exons whose inclusion is affected by transcription elongation rates and perturbations of RNA Pol II are among the 30% that appear to be cotranscriptionally excised (Naftelberg et al., 2015; Saldi et al., 2016; Herzel and Neugebauer, 2015).

Our data provide a rich resource for examining other questions of RNA metabolism and its regulation over development. Besides introns, transient species we can observe in the chromatin associated total RNA include upstream antisense RNAs and extended transcripts downstream from polyA sites (Seila et al., 2008; Flynn et al., 2011; Vilborg and Steitz, 2017) (data not shown). These data could also allow more sensitive detection of recursive or back-splicing, and inform studies of regulated RNA export. We are also examining regulated miRNA processing using additional data from short RNA libraries uploaded to GEO with the data discussed here (GEO(in process) and Supplemental Figure 2B) (Yeom et al., 2018).

### Behaviors of Retained Introns

To characterize incompletely spliced transcripts, we assessed introns based on their retention levels in different libraries, fractions and cell types. Unsupervised X-means clustering yielded four groups in each cell type. The largest cluster (Group A) were completely spliced in the poly(A)+ RNA, including in the chromatin fraction, and are presumably excised prior to transcription termination. The smallest cluster (Group D) behaved like classical retained introns in being exported to the cytoplasm within the otherwise fully spliced mRNA. Two intermediate clusters of introns (Groups B and C) were fully spliced in the cytoplasm while exhibiting different levels of retention on chromatin and to some extent the nucleoplasm. A deep neural network trained using a well-defined set of introns and a wide range of genomic features was able to distinguish introns in Group D from those in A or C with high accuracy. Group C introns were also distinguished from Group A with moderate accuracy (Figure 4B). These data indicate that Groups D and C are functionally distinct and the features which define them should give clues to their regulation. These features include those previously associated with retained introns, such as weak splice sites, conservation, and coding capacity (Sakabe and de Souza, 2007; Jaillon et al., 2008; Braunschweig et al., 2014; Dvinge and Bradley, 2015; Mauger et al., 2016; Parra et al., 2018), although we did not find a strong correlation between the presence of cryptic or decoy exons and intron retention, as described in other systems (Parra et al., 2018, 2020). We found that introns of the different groups were defined by enrichment of particular short sequence motifs in their terminal regions and adjacent exons. We have not yet identified proteins whose binding sites might underlie the enrichment of these motifs. This may be because the recognition elements assigned to individual proteins are not sufficiently specific. Alternatively, introns may be regulated by so many different proteins that no single binding motif is strongly predictive. Proteins including PTBP1 and others known to regulate particular retained introns demonstrate that such factors can be found (Yap et al., 2012; Horan et al., 2015; Frankiw et al., 2019b), but there may be many such factors each regulating a subset of introns in a group. The extension of our approach to larger datasets will allow correlation of changes in intron group assignment with the expression of particular RNA binding proteins.

Groups B and C include several previously described sets of interesting retained introns. Detained introns were defined as partially spliced introns in transcripts affecting growth control, whose excision can be modulated by cellular stimuli (Ninomiya et al., 2011; Boutz et al., 2015; Braun et al., 2017). These detained introns are a subset of the Group B and particularly Group C introns we defined in mESC. Another group of retained introns were shown to be regulated by PTB proteins in a neuronal cell line. Our analytical strategy identified many new PTBP1 dependent introns that remain as chromatin associated transcripts in mESC. Another study identified retained introns in the total cellular polyadenylated RNA of mature primary neuronal cultures (Mauger et al., 2016). This study defined retained introns whose splicing increased upon transcription inhibition as transient, and other introns that remained after transcriptional shutoff as stable. In our data from less mature neurons, we found that the largest group of transient introns were in Group C (40 %). In contrast, of the stable introns that we could assay in our cultures, about 40 % were in Group D (Supplemental Table S6E). This is consistent with the stable introns remaining in cytoplasmic mRNA after transcriptional shutoff. Similar to detained introns, Mauger et al. found that synaptic activation could change the splicing level of some retained introns. It will be interesting to examine whether these introns are associated with chromatin, but this will require improved isolation of nuclei from mature neuronal cultures.

### Developmental Regulation by Splicing Inhibition and Chromatin Sequestration

In previous studies, we showed how the neuronal specific expression of certain genes is determined by the coupling of a PTBP1 dependent splicing event to NMD. RNAs for the neuronal PTBP2 and PSD-95 proteins are expressed in ESC and other cells, but through the action of PTBP1 are spliced as isoforms that are subject to NMD (Boutz et al., 2007; Makeyev et al., 2007; Spellman et al., 2007; Zheng et al., 2012; Linares et al., 2015). A similar mechanism affects Gabbr1 through regulation of exon 15 by PTBP1 (Makeyev et al., 2007), but the change in RNA with loss of NMD is small compared to the complete absence of GABBR1 protein in ESC. Here we uncover another mechanism controlling the developmental specific expression of a neuronal protein. The Gabbr1 RNA is abundant in mESC but its splicing is incomplete and its transcript remains in the chromatin compartment.

Gabbr1 is expressed as multiple isoforms (Kaupmann et al., 1997). The long Gabbr1a isoform comes from a promoter active in all three cell types studied here. Gabbr1b, which lacks N-terminal sushi domains, arises from an alternative promoter within intron 5 active in neurons (Vigot et al., 2006). There is also a short transcript derived from an alternative polyA site in intron 4. A microexon 5 between these two introns adds a linker into the 1a isoform (Vigot et al., 2006). This complex intron 4-5 region is largely unprocessed in mESC cells, and becomes processed in neurons with the production of cytoplasmic mRNA including exon 5. The depletion of PTBP1 from mESC leads to multiple changes in Gabbr1 splicing including activation of microexon 5 and downstream exon 15. This leads to some cytoplasmic expression of neuronal mRNA isoforms but very limited protein expression. Much of the RNA remains nuclear indicating that additional factors prevent its mobilization. Instead of regulation at the level of transcription or mRNA stability, incomplete Gabbr1 splicing and sequestration of its RNA on chromatin are modulated to control gene output over development.

Most protein-coding transcripts exhibiting chromatin enrichment were not seen to be upregulated by Upf1 depletion. Other transcripts in this pool were similar to Gabbr1 in being modestly affected by NMD. The nuclear pools of these RNAs may reduce the observed efficiency of NMD on total RNA levels, where transcripts exhibit only partial depletion by the decay pathway even though near complete loss of protein is observed.

The Gabbr1 transcript is bound by PTBP1 across multiple sites. Studies have shown that when binding RNA at high stoichiometry, PTBP1 can cause the condensation of RNA/protein liquid droplets *in vitro*. It was recently found that extensive PTBP1 binding across repetitive binding sites in the long non-coding RNA Xist is required for Xist condensation onto the X chromosome during X inactivation (Pandya-Jones et al., 2020). PTBP1 also drives the condensation of the long non-coding RNA PNCTR in the perinucleolar compartment, and a similar mechanism may be involved in its interaction with LINE RNAs (Attig et al., 2018; Yap et al., 2018). It will be interesting to examine whether PTBP1 might create a nuclear condensate of Gabbr1 RNA. Although PTBP1 depletion led to increased splicing and increased mRNA in the soluble nucleoplasm and cytoplasm, it did not eliminate the enrichment of the unspliced RNA in the chromatin. This may be due to the partial depletion of PTBP1 by RNAi, but it seems likely that other proteins will also contribute to the sequestration of Gabbr1 RNA, as is seen with Xist. If the chromatin enrichment of protein-coding transcripts like Gabbr1 involve similar mechanisms to those controlling lncRNA function, they may also have similar effects on chromatin condensation and gene expression.

## MATERIALS AND METHODS

### Cell lines and tissue culture

The mESC line, E14 (Hooper et al. 1987) was cultured on 0.1 % gelatin-coated dishes with mitotically inactivated mouse embryonic fibroblasts (MEF) (CF1, Applied StemCell, Inc.) in mESC media at least 2 passages from the initial thawing. Then, the mESCs were transferred onto 100 mm cell culture plates that contained only 0.1% gelatin in preparation for RNAi and cell fractionation experiments. mESC media consisted of DMEM (Fisher Scientific) supplemented with 15 % ESC-qualified fetal bovine serum (Thermo Fisher Scientific), 1x non-essential amino acids (Thermo Fisher Scientific), 1x GlutaMAX (Thermo Fisher Scientific), 1x ESC-qualified nucleosides (EMD Millipore), 0.1 mM β-Mercaptoethanol (Sigma-Aldrich), and 10^3^ units/ml ESGRO leukemia inhibitor factor (LIF) (EMD Millipore). Mouse primary cortical neuron cultures were prepared from gestational day 15 C57BL/6 embryos (Charles River Laboratories), as described previously (Zheng 2010). Briefly, cortices were dissected out into ice cold HBSS and dissociated after a 10 min digestion in Trypsin (Thermo Fisher Scientific), then plated with plating media consisted of 70 % Neurobasal (v/v), 20 % horse serum (v/v), 25 mM sucrose, and 0.25x Glutamax plated at 5 million cells per 78.5 cm^2^ on 0.1 mg/ml poly-L-lysine coated dishes. The half media was replaced with feeding media consisted of 98% Neurobasal (v/v), 1x B27 (with vitamin A, Thermo Fisher Scientific), and 0.25x Glutamax at day 1 and 3. Neurons represented 70 – 80 % of the cells in the culture. A mouse neuronal progenitor cell line was established from cortical cells of gestational day 15 embryos generated by crossing homozygous Nestin-GFP transgenic mice to wild type C57BL/6. GFP positive cells were collected by FACS and plated on uncoated culture dishes. These NPCs were grown in DMEM/F12 supplemented with B27 (without vitamin A, Thermo Fisher Scientific), 1x GlutaMax and antibiotics. EGF and FGF (PeproTech) were added every day at 10 ng/ml concentration. All experiments were approved by the UCLA Institutional Animal Care and Use Committee (ARC# 1998-155-53).

### Knockdown of PTBP in mESC

siRNAs that target EGFP (Silencer Select AM4626), PTBP1 (Silencer Select s72337), and PTBP2 (Silencer Select s80149) were transfected into mESCs twice using Lipofectamine RNAiMAX (Invitrogen). The first transfection was performed while the cells were in suspension, and the second transfection was performed 24 hours after when the cells had already attached to the surface of the cell culture dish. The cells were harvested and fractionated 24 hours after the second transfection.

### Subcellular fractionation, RNA isolation, and library construction

Total RNA was isolated from mESCs, mNPC, and cortical neurons (mCtx) that were fractionated into cytoplasmic, soluble nuclear, and chromatin pellet compartments as described previously (Pandya-Jones 2009, Wuarin Schibler 1994, Yeom Damianov 2017, Yeom 2018). Briefly, cells were washed twice with washing buffer (1X PBS/1 mM EDTA [pH 8.0]) and gently trypsinized to collect the cell pellet by centrifugation. 1 × 10^7^ cells from the pellet were retrieved in a 2.0 mL low adhesion microcentrifuge tube (USA Scientific) and washed twice with washing buffer to remove cell debris from prematurely-lysed cells. For neurons, the initial plating number on the day of dissection and culture was used for quantification, and cells were collected by scraping from the plate without trypsinization.

The cells were then incubated first in ice-cold hypotonic buffer (10 mM Tris–HCl [pH 7.5], 15 mM KCl, 1 mM EDTA [pH 8.0], 0.15 mM Spermine, 0.5 mM Spermidine, 0.5 mM DTT, and 1× Protease inhibitor, and 15 mM KCl (50 mM NaCl for mNPC and mCtx)) for 5 minutes and lysed in ice-cold lysis buffer (10 mM Tris–HCl [pH 7.5], 15 mM KCl, 1 mM EDTA [pH 8.0], 0.15 mM Spermine, 0.5 mM Spermidine, 0.5 mM DTT, and 1× Protease inhibitor, and 0.0375%, 0.15%, and 0.1% of Igepal CA-630 for mESC, mNPC, and mCtx, respectively) for 1 minute. The resulting lysate was immediately layered on top of a chilled 24 % sucrose solution (hypotonic buffer in 24 % (w/v) sucrose without detergent) and centrifuged for 10 minutes, 4 °C, 6000 x g. 10% of the supernatant (cytoplasmic fraction) was used for immunoblot to check for contamination from nuclear materials, and the rest was mixed with Trizol LS (Invitrogen) to extract cytoplasmic RNA. The nuclear pellet was washed once with washing buffer before being resuspended in 100 μl of chilled glycerol buffer (20mM Tris-HCl [pH 7.9], 75 mM NaCl, 0.5 mM EDTA [pH 8.0], 50% glycerol (v/v), 0.85 mM DTT, and 0.125 mM PMSF). 100 μl of cold nuclear lysis buffer (20 mM HEPES [pH 7.6], 7.5 mM MgCl2, 0.2 mM EDTA [pH 8.0], 300 mM NaCl, 1M urea, 1% Igepal CA-630, 0.3 mM Spermine, 1.0 mM Spermidine, 1× Protease inhibitor) was then added and the mixture incubated on ice for 2 minutes. After the 2 minutes incubation period, 1/8 of the nuclear lysate was transferred to a separate 1.5 mL microcentrifuge tube for each sample. After centrifugation for 2 minutes, 4 °C, 6000 x g, the supernatant (soluble nuclear fraction) from both tubes were pooled. 10% of the pooled supernatant was used to check its purity by western blot and the rest was mixed with Trizol LS to extract nucleoplasmic RNA. The resulting two insoluble nuclear pellets (one large, one small) were washed once with washing buffer before incubation of the large pellet in Trizol at 50 °C until the pellet was completely solubilized to extract chromatin-associated RNA (from 7/8 nuclear lysate). After washing the small pellet (from 1/8 nuclear lysate) was resuspended in 5% SDS sample buffer (5% SDS, 30% glycerol (v/v) and 150 mM Tris–HCl [pH 8.0]) at 55 °C for 15 minutes to extract proteins from the chromatin fraction. Each fractionation experiment was performed in triplicate.

RNA was treated DNase I (Takara) followed by phenol extraction. Before RNA quality check and the size selection steps, RNA was quantified on a nanodrop spectrophotomer (Supplemental Table S2A), and RNA integrity was checked by either Bioanalyzer or RNA screen tape (Agilent) (Supplemental Table S2A). Note that the sum of the RNA from the chromatin, nucleus, and cytoplasmic fractions does not represent total cellular RNA due to extensive washing steps during fractionation.

5ug of RNA from each subcellular fraction was processed with the RNeasy MinElute Cleanup Kit (Qiagen). RNAs longer than 200 nt were eluted from the membrane to make long RNA libraries, while the flow through (< 200 nt) was also collected and ethanol precipitated to make short RNA libraries from each fraction. Half of the long RNA eluate was used to generate poly(A)+ libraries using the TruSeq Stranded mRNA Library Prep Kit (Illumina). The other half the long RNA eluate was used for total RNA library construction, using the ribosomal RNA removal kit (Ribo-Zero™ rRNA Removal Kits (Human/Mouse/Rat), MRZH116, Epicentre Biotechnologies) followed by library construction using TruSeq Stranded mRNA Library Prep Kit (Illumina) without the oligo-T bead purification step. Short RNA was used for small RNA library construction, using the ribosomal RNA removal kit (Ribo-Zero™ rRNA Removal Kits (Human/Mouse/Rat), MRZH116, Epicentre Biotechnologies) followed by library construction using the TruSeq Small RNA sample Prep Kit (Illumina). The small RNA library was further purified on a TBE gel by excising bands between 120 - 160 nt to generate small non-coding RNA reads that include mature-miRNAs (~ 22 nt) and piwi-interacting RNAs (piRNAs, ~ 30 nt).

For PTBP knockdown experiments in mESC, poly(A)+ libraries were created using the TruSeq Stranded mRNA Library Prep Kit (Illumina).

### RNA sequencing and alignment

The libraries of all fractions and cell types, were subjected to 100 nt paired-end sequencing at the UCLA Broad Stem Cell Center core facility on an Illumina HiSeq4000 to generate about 21 million mapped reads per sample. The libraries of PTBP1 KD samples in mESC were subjected to 75 nt paired-end sequencing at the UCLA Neuroscience Genomics Core on an Illumina HiSeq4000 to generate about 25 million mapped reads per sample. RNA-seq data were aligned using STAR (version 020210) (Dobin et al., 2013) on mm10. Small RNA libraries were sequenced on an Illumina MiSeq machine using the MiSeq Reagent Kit v3 (Illumina).

Sequencing reads and mapping results are summarized in Supplemental Table S1B – S1D. Reads per million read values (RPM) were calculated and displayed in UCSC genome browser sessions including strand information. TPM values obtained from Kallisto (version 0.43.0) were used in gene expression analyses (Bray et al., 2016). To examine the similarity of replicate samples, gene expression distances between samples were calculated using DESeq2 (Anders and Huber, 2010), and plotted in heatmaps (Supplemental Figure S1) using ggplot2.

### Accession numbers and genome browser sessions

All sequence data was deposited in GEO under the accession number [GEO:in process]. Links to the data displayed on the UCSC genome browser here (https://genome.ucsc.edu/s/Chiaho/Kay_fraction_total_hub_10202020 for total RNA, and https://genome.ucsc.edu/s/Chiaho/Kay_fraction_polyA%2B_hub_10202020 for poly(A)+ RNA)

### Calculation of chromatin partition indices and biotype analysis

To analyze differential compartmentalization of RNAs, genes were selected to pass the following expression filters: having chromatin expression greater or equal to the median TPM reported by Kallisto (2.13 TPM), and having read counts greater than 0 in the cytoplasmic fractions as measured by FeatureCount. This returned 13,036 genes for analysis. DEseq2 was used to measure fold change in read counts between the chromatin-associated and the cytoplasmic poly(A)+ RNA by calculating the average read count between replicates of the chromatin fraction divided by the average read counts of the cytoplasmic fraction. The chromatin partition index was defined as the log2 transformation of this ratio (Figure 1D).

Biotype information was retrieved from Ensembl annotation (V.91). Of the total 13,036 genes, 400 genes (3.1%) were analyzed in each of three ranges of the distribution. Partition indices were from −4.2 to −2.6 for region L, −0.1 to 0.1 for region M, and 4.1 to 8.6 for region R.

### Co- and posttranscriptional splicing analysis

For cotranscriptional splicing analysis, we filtered the U intron list for genes whose Kallisto TPM in chromatin poly(A)+ RNA was over the median. We then identified genes exhibiting 3’ bias by comparing the reads per nucleotide length from the 2^nd^ exon to the reads per nucleotide length of the 2^nd^ to last exon for the longest transcript from the gene. If the ratio of 2^nd^ exon reads to 2^nd^ to last exon reads was less than 0.5 for all transcripts, we excluded that gene from the analysis. The following steps were described in Supplemental Figure S3C. These filters were also applied to a list of simple cassette exons (SE) extracted from rMATS 4.0.1 (Shen et al., 2014). The subset of these cassette exons was selected that are located between I introns when they included, and become EI introns when skipped as defined in Supplemental Figure S3E.

To examine splicing at different intron locations, we compiled a list of 936 genes containing U introns as first, middle, and last introns. The middle intron in these genes was defined as the intron number X, depending on whether the gene has an even or odd number of total introns. For an odd number of total introns, X= (total intron number + 1) / 2. If the gene has an even number of total introns, then X= total intron number/2.

### RT-PCR and RT-qPCR validation

RNA was collected and extracted from subcellular fractions as described above. For the RT reaction, 0.8 - 1 μg of total RNA, 100 ng of random hexamers (or 250 ng of oligo dT), and 0.5 mM of dNTP mix (0.5 mM each) were incubated in a 7 μl reaction volume at 65 °C for 5 min. After 1min incubation on ice, 2 μl of 5x reaction buffer, 5 mM of DTT, and 100 units of SuperScript III RT (Thermo Fisher Scientific) were added. This 10 μl mixture was incubated at 25 °C for 10 min, 50 °C for 60 min, and 70 °C for 15 min. PCR was performed using Phusion DNA polymerase (Fisher Scientific). After the initial denaturing step at 98 °C for 1min, amplification continued for 19 - 28 cycles of 98 °C for 1min, 58 - 62 °C for 30 sec (annealing), and 72 °C for 20 - 25 sec (elongation). The reaction was completed with a final elongation at 72 °C for 10min. Annealing temperature, elongation time, cycle numbers, and amplicon size are listed in Supplemental Table S1. RT-PCR products were run on either agarose gels with EtBr staining or PAGE gels with SYBR Gold staining (Thermo Fisher Scientific), and visualized on a Typhoon imager (GE Healthcare) using the 492 nm excitation laser and 510 nm emission filters. For RT-qPCR the SensiFAST SYBR Lo-ROX Kit (Bioline) was used, and reactions contained 0.4ul of diluted cDNA (1:5 dilution in water) with 250nM each of forward and reverse primers in 6ul of RT-qPCR reactions. The mixtures were run on a QuantStudio 6 Real-Time PCR System (Thermo Fisher Scientific) with combined annealing and elongation cycles (Step1: 95 °C for 2min, followed by 40 cycles of Step2: 95 °C for 5sec and Step3: 55 or 60°C for 30 sec. A list of primer sequences and PCR conditions is presented in Supplemental Table S1.

### Measurement of Intron Retention

We developed SIRI (Systematic Investigation of Retained Introns), a tool to stringently quantify unspliced introns by deep sequencing (https://github.com/Xinglab/siri). In this tool, we first retrieve all introns from Ensembl gene transfer format (GTF) version 91 for the mouse mm10 genome (citation). Then we count the number of reads that map to exon-exon (EE), exon-intron (EI), and intron-exon (IE) junctions. The FI (Fraction of intron Inclusion) value indicates the fraction of the intron retained in the transcripts. We selected only U type IR events from poly(A)+ selected RNA-seq experiments. We required that the IR events have an intron length greater than and equal to 60, and that the sum of EE + EI + IE reads be greater than and equal to 20 (https://xinglabtrackhub.research.chop.edu/intron_retention/ for U introns, https://xinglabtrackhub.research.chop.edu/intron_retention/TS7_Intron_FI_combined.xls x for all types of introns). From this set, IR events with EE reads no fewer than 2 in at least one cell compartment in one cell type were then kept for downstream analysis.

### X-means Clustering of IR events

We applied X-means clustering to the FI values determined in all three compartments of each cell type. X-means does not require a user-defined cluster number. Instead, it automatically determines the cluster number by optimizing the Bayesian Information Criteria (BIC). We used the pyclustering tool (Novikov, 2019) for the clustering analysis with the maximum number of clusters set at 6. The distance matrix for X-means clustering is based on the Dynamic Time Warping (DTW) algorithm (Berndt and Clifford, 1994) for the purpose of investigating directional changes of FI values from chromatin to nucleoplasm to cytoplasm. The circos plot showing the intron group changes from one cell type to another cell type was produced using R package circlize (version 0.4.4).

### Predicting intron retention patterns by deep learning

To apply deep learning to the prediction of IR groups, we first constructed a compendium of 1,387 intron features of five types: sequence motifs, transcript features, RNA secondary structure, nucleosome positioning, and conservation (Supplemental Table S7). Sequence motif features included splice site consensus sequences, position-specific matrices of RNA binding proteins, dinucleotide and trinucleotide frequencies of introns and flanking exons. Transcript features include the lengths of upstream exon (E1), downstream exon (E2), and intron (I) and intron number in its host gene. The translatability of E1, E2, E1+E2, I and E1 + I + E2 were defined by checking whether there was stop codon in at least one of three possible reading frames. To predict RNA secondary structure, RNA sequences from the regions of 5’ of exon-intron boundary (−20nt into exon and +20nt into intron), 3’ of intron-exon boundary (−20nt into intron and +20nt into exon) were examined. Similarly, sequence intervals from 1 to 70nt, 70 to 140nt, 140 to 210nt from the 5’ portion of the intron, and from −210 to −140nt, −140 to −70nt and −70nt to −1nt from the 3’ portion of the intron were used. We computed the free energy of folding for each region with RNAfold (2.2.10) (Kerpedjiev P et al. 2015, Bioinformatics) and the free energy of unfolding for each region was used as a feature for the deep learning. The nucleosome positioning was predicted by NuPoP (version 1.0, set to the mouse model) on the last 50nt of the upstream exon, the first 100nt of 5’ intron region, the last 100nt of 3’ intron region, and the first 50nt of downstream exon [citation]. We collected a reliable training dataset by including introns that had grouping information in at least two cell types and removing U11/U12 introns and other introns lacking GT or AG splice sites. We trained a Deep Neural Network (DNN) with these 1,387 features to predict whether introns belong to group A, B, C, and D for each cell type (Figure 3A). The training was done with five-cross validation and the area under the ROC (Receiver Operating Characteristic) curve on data held-out during training was reported for performance evaluation.

### Identifying Premature Termination Codons (PTC) in retained introns

For each U intron, we retrieved all transcripts containing this intron from GTF. The transcripts without a clear open reading frame annotation or with termination codons after the intron were not considered. We also removed transcripts in which the intron was the last intron. Because last intron retention should not cause Nonsense Mediated Decay (NMD). For each transcript that contained the intron, we inserted the intron sequence into the CDS of the transcript at junction of the intron. If the U intron was not the second last intron of the transcript, the intron was marked as PTC-containing if generated a stop codon inside the intron. If the intron was the second last intron of the transcript, the intron was defined as PTC-containing if an in frame stop codon within the intron was at least 50nt upstream from the last exon-exon junction of the transcript. After checking all transcripts containing the intron, we defined the intron as a PTC-containing intron if it generated PTC in at least one transcript.

### GO analysis

Gene ontology enrichments were determined using PANTHER (Protein ANalysis THrough Evolutionary Relationships) (Ashburner et al., 2000; Gene Ontology Consortium, 2019). Molecular function terms were assessed for overrepresentation compared to all mouse genes in the list of genes containing introns that changed during neuronal differentiation from Groups C or D to Groups A or B, or from Groups A or B to Groups C or D.

### Identification of NMD target genes

A list NMD targets was extracted from a previous study in mESC (Hurt et al., 2013). We filtered for genes upregulated by greater than 10% in two independent siRNA knockdowns of Upf1 (siUpf1.1 and siUpf1.2). If a gene had multiple mRNA isoforms and one isoform was upregulated it was included as an NMD target on our list.

## ACKNOWLEDGEMENTS

We thank Grigori Enikolopov (Cold Spring Harbor Laboratory) for the Nestin-GFP mouse line; Celine Vuong for help with neuronal culture; and Amy Pandya-Jones, Kathrin Plath, and members of the Black lab for discussions and comments on the manuscript. This work was supported by a W.M. Keck Foundation grant and NIH grant R01 MH109166 to DLB and YX, NIH grants R35 GM136426 and R01 GM049662 to DLB, R01 GM088342 to YX, and support from the David Geffen School of Medicine and Division of Life Sciences at University of California, Los Angeles (UCLA) to DLB. K-HY was supported by a postdoctoral fellowship from Eli and Edythe Broad Center of Regenerative Medicine and Stem Cell Research at UCLA. HL was supported by Whitcome Fellowship from the UCLA Molecular Biology Institute and a Warsaw Family graduate fellowship from the department of MIMG from the UCLA.

---

**Supplemental Figure S1. (Related to Figure 1).**
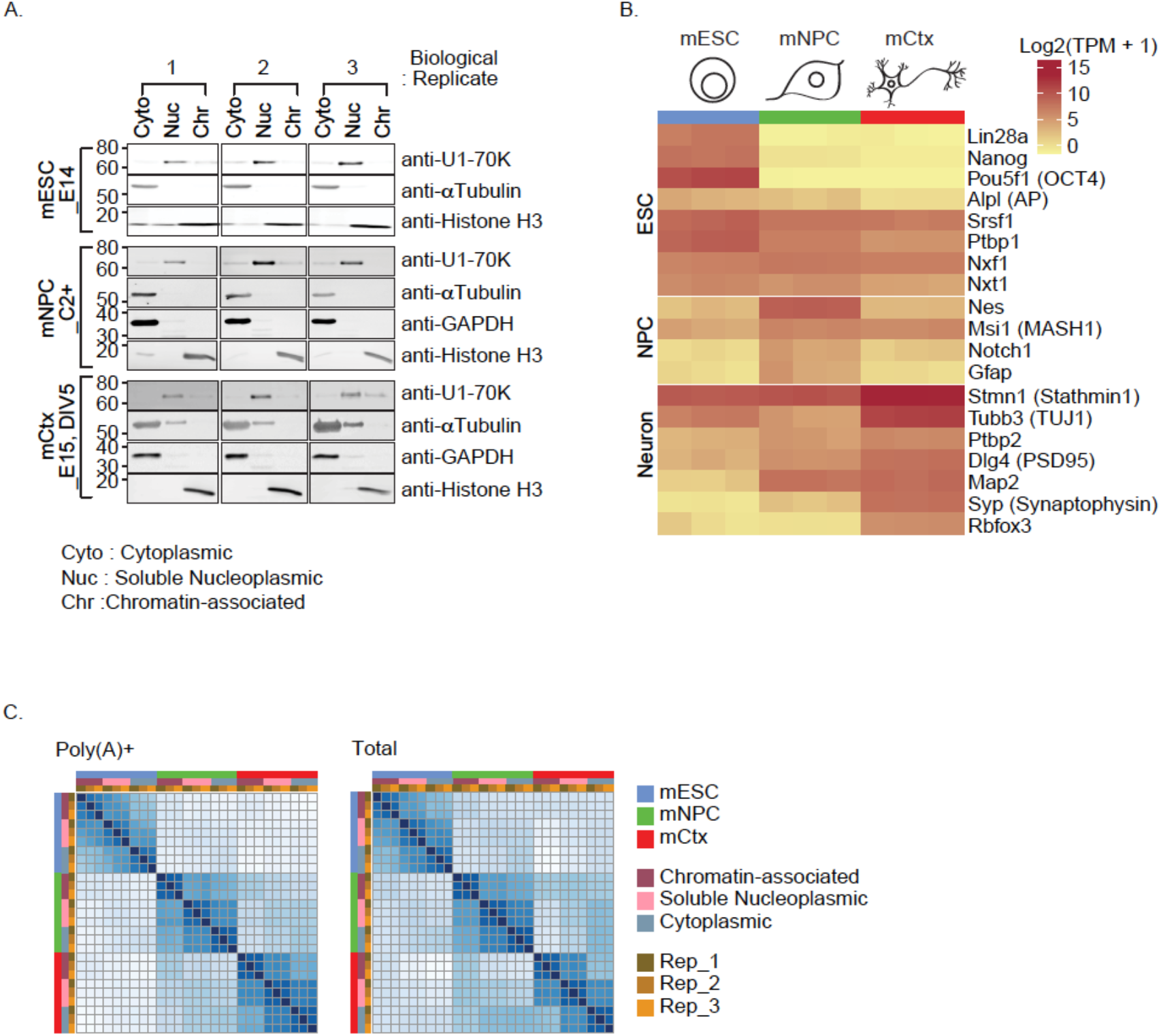
Validation of subcellular fractionation, cell type gene expression, and library consistency. A. Confirmation of subcellular fractionation. Immunoblot analysis of diagnostic proteins in sub cellular fractions. U1-70K for soluble nucleoplasm (Nuc), alpha-TUBULIN and GAPDH for cytoplasm (Cyto), and HISTONE H3 for chromatin pellet (Chr). Gel images include 3 biological replicates of mouse embryonic stem cells (line E14), mouse neuronal progenitor cell line C2+, and mouse cortical neurons after 5 days *in vitro* culture (mCtx). Note that the immunoblot results of the third replicate of mESC_E14 was published previously (Yeom et al., 2018). B. Confirmation of cell type specific gene expression. Heatmap presents the cytoplasmic expression as measured by Kallisto for the indicated mRNAs in each cell type and replicate. C. Confirmation of library similarity. Heatmap displays similarity of gene expression between pairwise comparisons of all cell types, fractions, and replicates. Color codes are indicated on right.

**Supplemental Figure S2. (Related to Figure 1).**
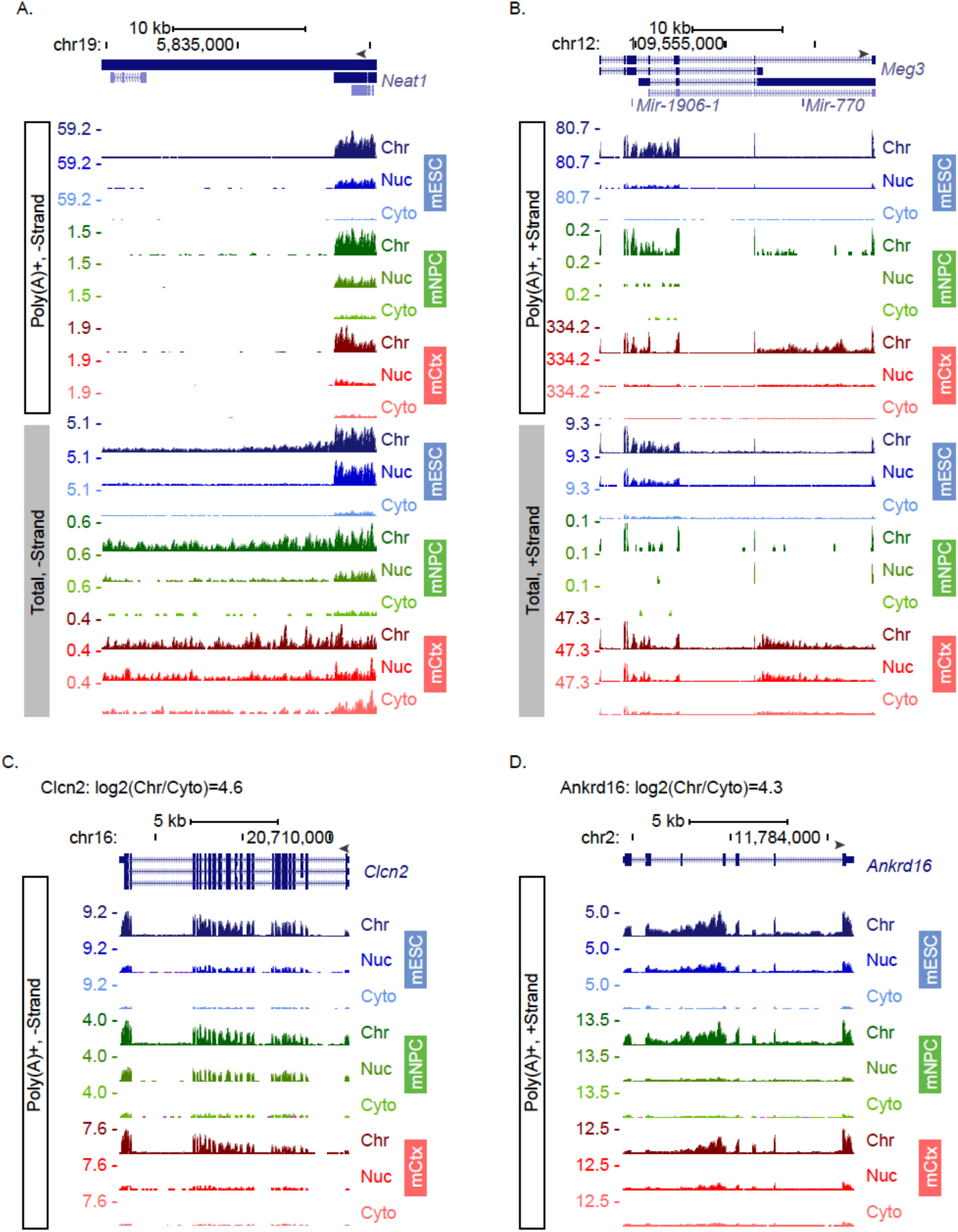
Example genome browser tracks of non-coding and coding RNAs. A. Neat1 expression in mESC, mNPC, and mCtx. Genome browser tracks of the *Neat1* locus for poly(A)+ and Total libraries. Y axis shows RPM scaled to the highest value in the Chromatin-associated fraction. B. Meg3 expression in mESC, mNPC, and mCtx. Genome browser tracks of the *Meg3* locus in poly(A)+ and Total libraries. C. Genome browser tracks of the *Clcn2* locus in mESC, mNPC, and mCtx. Transcripts are enriched in the chromatin fraction and exhibit unspliced introns in poly(A)+ RNA. D. Genome browser tracks of the *Ankrd16* locus in mESC, mNPC, and mCtx. Transcripts are enriched in the chromatin fraction and exhibit unspliced introns in poly(A)+ RNA.

**Supplemental Figure S3. (Related to Figures 2 and 3).**
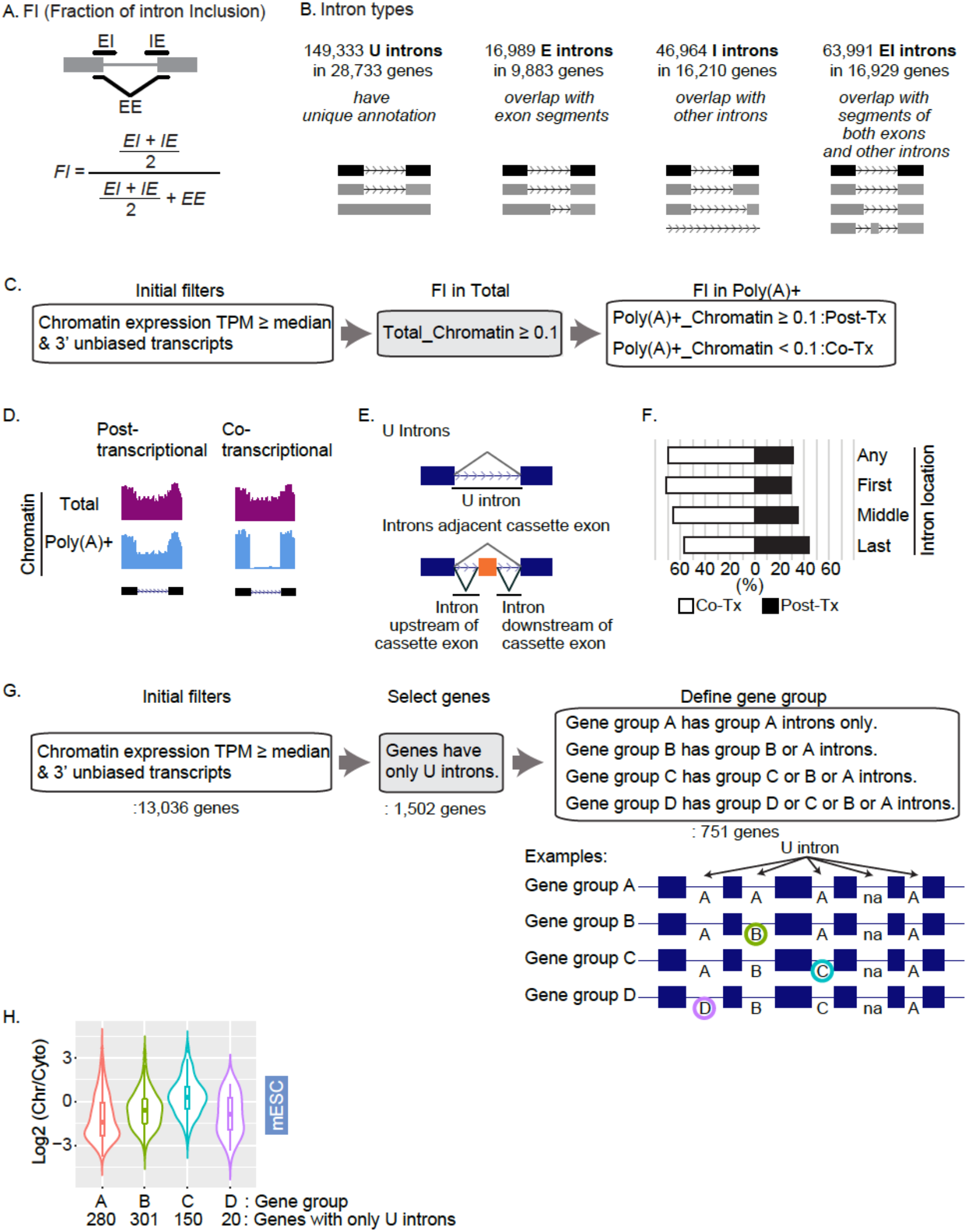
Computational definition of introns and splicing. A. Determination of FI value using read numbers for the exon-intron junction (EI), intron-exon junction (IE), and exon-exon junction (EE). B. Introns were categorized as one of four types based on their Ensembl v91 annotation. Introns that are not partly overlapped with either exons or other introns are classified as U type introns. Introns that partly overlap with exons but not with other introns are classified E type introns. Introns that overlap with other annotated introns but not exons are called I type introns. EI type introns overlap with both exons and introns of other annotated isoforms. C. Determination of cotranscriptional and posttranscriptional splicing. FI values were determined for all U introns from total and poly(A)+ chromatin associated RNA. Genes with overall expression above the median (2.13 TPM) were analyzed. Genes showing a bias for reads in the 3’ end in the poly(A)+ RNA, and introns exhibiting FI values in total RNA below 0.1 were removed. A posttranscriptional splicing event was then defined as an intron having an FI value in Poly(A)+ RNA greater than or equal to 0.1 (Post-tx). Cotranscriptional splicing of an intron generates an FI of less than 0.1 in the poly(A)+ RNA (Co-tx). D. Illustration of post and cotranscriptional splicing. Introns with high read numbers on chromatin in both the total and poly(A)+ libraries were defined as posttranscriptionally spliced. Cotranscriptional splicing events exhibited reads in the total but not the poly(A)+ RNA. E. Diagrams of constitutive U introns and I introns adjacent to simple cassette exons that were assessed for co-and posttranscription splicing as presented in Figure 2C. F. The proportions of co- and posttranscriptional splicing for all U introns and for first, middle and last introns in a transcript. G. Transcripts with unspliced introns are enriched in the chromatin fraction. Genes having only U introns were selected from those whose overall expression was above the median (2.13 TPM). The gene group was then defined by the highest intron group within the gene (751 genes), where D>C>B>A. Introns marked ‘na’ indicate they were filtered by SIRI during X-means clustering. H. Violin plots showing the distribution of chromatin partition indices (Log2(Chr/Cyto)) of transcripts from different gene groups defined above. The number of genes in each gene group is indicated at the bottom.

**Supplemental Figure S4. (Related to Figure 5).**
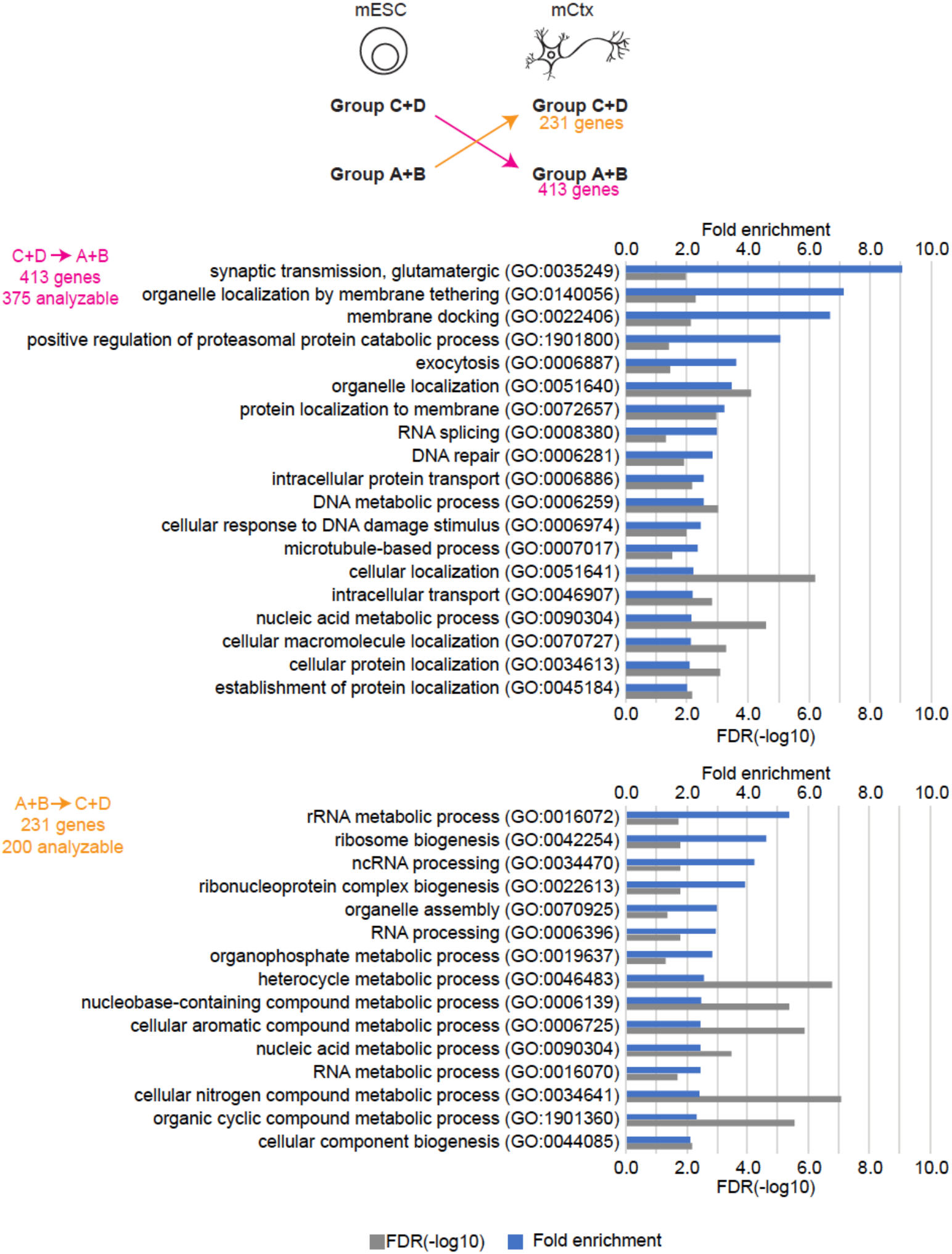
GO analysis of genes containing introns that switch intron group during neuronal differentiation. Number of genes containing introns that changed group between mESC and mCtx is indicated at the top in yellow and pink. GO biological process enrichment these gene sets are listed at the bottom. Fold enrichment and FDR (−log10) shown in blue and grey bars, respectively.

**Supplemental Figure S5. (Related to Figure 5 and Figure 6).**
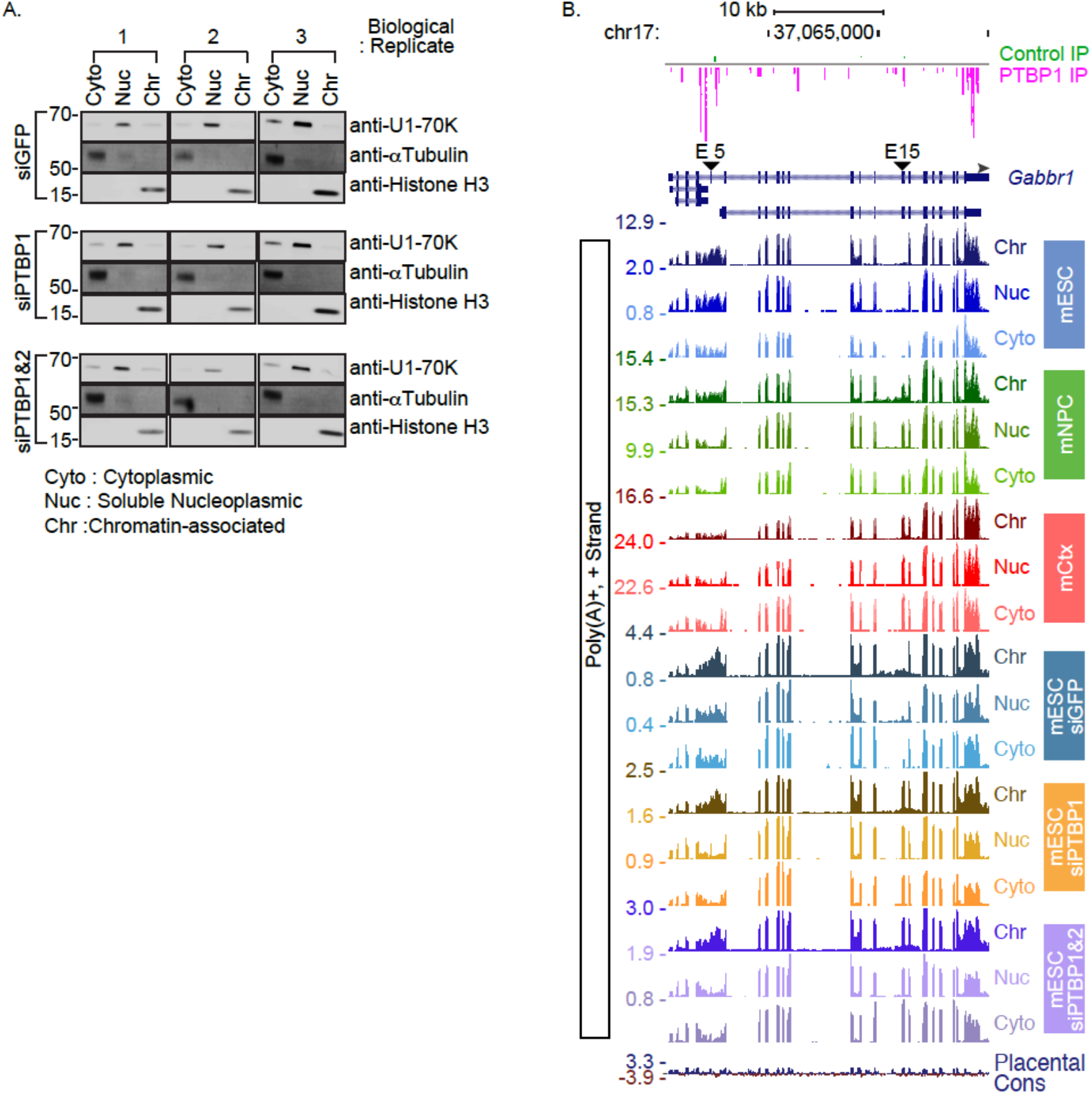
Validation of subcellular fractionation after PTBP knockdown in mESC and genome browser tracks of Gabbr1. A. Confirmation of subcellular fractionation. Immunoblot analysis of diagnostic proteins in sub cellular fractions. U1-70K for soluble nucleoplasm (Nuc), alpha-TUBULIN and GAPDH for cytoplasm (Cyto), and HISTONE H3 for chromatin pellet (Chr). Gel images include 3 biological replicates of mouse embryonic stem cells (line E14). B. Complete genome browser tracks of the *Gabbr1* locus in mESC, mNPC, and mCtx, and for PTBP1 knockdown and PTBP1/2 double knockdown in mESC. PTBP1 iCLIP tags in mESC are shown at the top. Intron 4-5 region is shown with a bracket, and exons 5 and 15 are marked with arrowheads.

## SUPPLEMENTAL TABLES

**Supplemental Table S1.**

Primers, siRNAs and Antibodies used in this study, and PCR conditions.

**Supplemental Table S2.**

(A) Quantification of RNA extracted from subcellular fractions and (B-D) summary of genome alignment statistics.

**Supplemental Table S3.**

Comparison of RNA-seq RPM and qPCR quantification between subcellular fractions.

**Supplemental Table S4.**

Analyses of co- and posttranscriptional splicing (related to Figure 2C).

**Supplemental Table S5.**

Properties of retained introns.

(A) Introns containing premature termination codons (Related to Figure 3C) and (B) proportions of C and D introns located in UTRs (Related to Figures 3D and 4C).

**Supplemental Table S6.**

Lists of introns and genes from previous studies that were analyzed in this study.

(A and B) Lists of detained introns from Boutz et al. (2015) analyzed as U introns in this study (Related to Figure 3D). (C) List of genes containing PTBP1 dependent retained introns in chromatin fraction that were upregulated after Upf depletion and (D) summary of genes with different chromatin partition indices in mESC that were up regulated after Upf1 knock down (Hurt et al., 2013). (E) Overlap of U introns and intron groups in mCtx that overlap with intron groups from Mauger et al. (2016).

**Supplemental Table S7.**

List of sequence features for the deep leaning model, and predictive value of genomic features for distinguishing introns of different groups.

## REFERENCES

Ameur, A., Zaghlool, A., Halvardson, J., Wetterbom, A., Gyllensten, U., Cavelier, L., and Feuk, L. (2011). Total RNA sequencing reveals nascent transcription and widespread co-transcriptional splicing in the human brain. Nat. Struct. Mol. Biol. 18, 1435–1440.

Anders, S., and Huber, W. (2010). Differential expression analysis for sequence count data. Genome Biol. 11, R106.

Ashburner, M., Ball, C.A., Blake, J.A., Botstein, D., Butler, H., Cherry, J.M., Davis, A.P., Dolinski, K., Dwight, S.S., Eppig, J.T., et al. (2000). Gene Ontology: tool for the unification of biology. Nat. Genet. 25, 25–29.

Attig, J., Agostini, F., Gooding, C., Chakrabarti, A.M., Singh, A., Haberman, N., Zagalak, J.A., Emmett, W., Smith, C.W.J., Luscombe, N.M., et al. (2018). Heteromeric RNP Assembly at LINEs Controls Lineage-Specific RNA Processing. Cell 174, 1067–1081.e17.

Berndt, D.J., and Clifford, J. (1994). Using dynamic time warping to find patterns in time series. In Proceedings of the 3rd International Conference on Knowledge Discovery and Data Mining, (Seattle, WA: AAAI Press), pp. 359–370.

Bhatt, D.M., Pandya-Jones, A., Tong, A.-J., Barozzi, I., Lissner, M.M., Natoli, G., Black, D.L., and Smale, S.T. (2012). Transcript Dynamics of Proinflammatory Genes Revealed by Sequence Analysis of Subcellular RNA Fractions. Cell 150, 279–290.

Boutz, P.L., Stoilov, P., Li, Q., Lin, C.-H., Chawla, G., Ostrow, K., Shiue, L., Ares, M., and Black, D.L. (2007). A post-transcriptional regulatory switch in polypyrimidine tract-binding proteins reprograms alternative splicing in developing neurons. Genes Dev. 21, 1636–1652.

Boutz, P.L., Bhutkar, A., and Sharp, P.A. (2015). Detained introns are a novel, widespread class of post-transcriptionally spliced introns. Genes Dev. 29, 63–80.

Braun, C.J., Stanciu, M., Boutz, P.L., Patterson, J.C., Calligaris, D., Higuchi, F., Neupane, R., Fenoglio, S., Cahill, D.P., Wakimoto, H., et al. (2017). Coordinated Splicing of Regulatory Detained Introns within Oncogenic Transcripts Creates an Exploitable Vulnerability in Malignant Glioma. Cancer Cell 32, 411–426.e11.

Braunschweig, U., Barbosa-Morais, N.L., Pan, Q., Nachman, E.N., Alipanahi, B. Gonatopoulos-Pournatzis, T., Frey, B., Irimia, M., and Blencowe, B.J. (2014). Widespread intron retention in mammals functionally tunes transcriptomes. Genome Res. 24, 1774–1786.

Bray, N.L., Pimentel, H., Melsted, P., and Pachter, L. (2016). Near-optimal probabilistic RNA-seq quantification. Nat. Biotechnol. 34, 525–527.

Brockdorff, N., McCabe, M., Norris, P., Cooper, J., Swift, S., and Kay, F. (1992). The Product of the Mouse Xist Gene Is a 15 kb Inactive X-Specific Transcript Containing No Consewed ORF and Located in the Nucleus. 12.

Broseus, L., and Ritchie, W. (2020). Challenges in detecting and quantifying intron retention from next generation sequencing data. Comput. Struct. Biotechnol. J. 18, 501–508.

Coulon, A., Ferguson, M.L., de Turris, V., Palangat, M., Chow, C.C., and Larson, D.R. (2014). Kinetic competition during the transcription cycle results in stochastic RNA processing. ELife 3, e03939.

Davidson, L., Kerr, A., and West, S. (2012). Co-transcriptional degradation of aberrant pre-mRNA by Xrn2. EMBO J. 31, 2566–2578.

Dobin, A., Davis, C.A., Schlesinger, F., Drenkow, J., Zaleski, C., Jha, S., Batut, P., Chaisson, M., and Gingeras, T.R. (2013). STAR: ultrafast universal RNA-seq aligner. Bioinformatics 29, 15–21.

Duff, M.O., Olson, S., Wei, X., Garrett, S.C., Osman, A., Bolisetty, M., Plocik, A., Celniker, S.E., and Graveley, B.R. (2015). Genome-wide identification of zero nucleotide recursive splicing in Drosophila. Nature 521, 376–379.

Dvinge, H., and Bradley, R.K. (2015). Widespread intron retention diversifies most cancer transcriptomes. Genome Med. 7, 45.

Edfors, F., Danielsson, F., Hallström, B.M., Käll, L., Lundberg, E., Pontén, F., Forsström, B., and Uhlén, M. (2016). Gene-specific correlation of RNA and protein levels in human cells and tissues. Mol. Syst. Biol. 12, 883.

Fei, J., Jadaliha, M., Harmon, T.S., Li, I.T.S., Hua, B., Hao, Q., Holehouse, A.S., Reyer, M., Sun, Q., Freier, S.M., et al. (2017). Quantitative analysis of multilayer organization of proteins and RNA in nuclear speckles at super resolution. J. Cell Sci. 130, 4180–4192.

Flynn, R.A., Almada, A.E., Zamudio, J.R., and Sharp, P.A. (2011). Antisense RNA polymerase II divergent transcripts are P-TEFb dependent and substrates for the RNA exosome. Proc. Natl. Acad. Sci. 108, 10460–10465.

Frankiw, L., Baltimore, D., and Li, G. (2019a). Alternative mRNA splicing in cancer immunotherapy. Nat. Rev. Immunol. 19, 675–687.

Frankiw, L., Majumdar, D., Burns, C., Vlach, L., Moradian, A., Sweredoski, M.J., and Baltimore, D. (2019b). BUD13 Promotes a Type I Interferon Response by Countering Intron Retention in Irf7. Mol. Cell 73, 803–814.e6.

Garland, W., and Jensen, T.H. (2020). Nuclear sorting of RNA. WIREs RNA 11, e1572.

Gene Ontology Consortium, T. (2019). The Gene Ontology Resource: 20 years and still GOing strong. Nucleic Acids Res. 47, D330–D338.

Girard, C., Will, C.L., Peng, J., Makarov, E.M., Kastner, B., Lemm, I., Urlaub, H., Hartmuth, K., and Lührmann, R. (2012). Post-transcriptional spliceosomes are retained in nuclear speckles until splicing completion. Nat. Commun. 3, 994.

Hao, S., and Baltimore, D. (2013). RNA splicing regulates the temporal order of TNF-induced gene expression. Proc. Natl. Acad. Sci. 110, 11934–11939.

Hautbergue, G.M. (2017). RNA Nuclear Export: From Neurological Disorders to Cancer. In Personalised Medicine: Lessons from Neurodegeneration to Cancer, S. El-Khamisy, ed. (Cham: Springer International Publishing), pp. 89–109.

Herzel, L., and Neugebauer, K.M. (2015). Quantification of co-transcriptional splicing from RNA-Seq data. Methods 85, 36–43.

Herzel, L., Ottoz, D.S.M., Alpert, T., and Neugebauer, K.M. (2017). Splicing and transcription touch base: co-transcriptional spliceosome assembly and function. Nat. Rev. Mol. Cell Biol. 18, 637–650.

Horan, L., Yasuhara, J.C., Kohlstaedt, L.A., and Rio, D.C. (2015). Biochemical identification of new proteins involved in splicing repression at the Drosophila P-element exonic splicing silencer. Genes Dev. 29, 2298–2311.

Hurt, J.A., Robertson, A.D., and Burge, C.B. (2013). Global analyses of UPF1 binding and function reveal expanded scope of nonsense-mediated mRNA decay. Genome Res. 23, 1636–1650.

Hutchinson, J.N., Ensminger, A.W., Clemson, C.M., Lynch, C.R., Lawrence, J.B., and Chess, A. (2007). A screen for nuclear transcripts identifies two linked noncoding RNAs associated with SC35 splicing domains. BMC Genomics 8, 39.

Jacob, A.G., and Smith, C.W.J. (2017). Intron retention as a component of regulated gene expression programs. Hum. Genet. 136, 1043–1057.

Jaillon, O., Bouhouche, K., Gout, J.-F., Aury, J.-M., Noel, B., Saudemont, B., Nowacki, M., Serrano, V., Porcel, B.M., Ségurens, B., et al. (2008). Translational control of intron splicing in eukaryotes. Nature 451, 359–362.

Kaupmann, K., Huggel, K., Heid, J., Flor, P.J., Bischoff, S., Mickel, S.J., McMaster, G., Angst, C., Bittiger, H., Froestl, W., et al. (1997). Expression cloning of GABA B receptors uncovers similarity to metabotropic glutamate receptors. Nature 386, 239–246.

Keppetipola, N., Sharma, S., Li, Q., and Black, D.L. (2012). Neuronal regulation of pre-mRNA splicing by polypyrimidine tract binding proteins, PTBP1 and PTBP2. Crit. Rev. Biochem. Mol. Biol. 47, 360–378.

Khodor, Y.L., Menet, J.S., Tolan, M., and Rosbash, M. (2012). Cotranscriptional splicing efficiency differs dramatically between Drosophila and mouse. RNA 18, 2174–2186.

Li, Y., Bor, Y.-C., Misawa, Y., Xue, Y., Rekosh, D., and Hammarskjöld, M.-L. (2006). An intron with a constitutive transport element is retained in a Tap messenger RNA. Nature 443, 234–237.

Linares, A.J., Lin, C.-H., Damianov, A., Adams, K.L., Novitch, B.G., and Black, D.L. (2015). The splicing regulator PTBP1 controls the activity of the transcription factor Pbx1 during neuronal differentiation. ELife 4, e09268.

Liu, H., Liang, C., Kollipara, R.K., Matsui, M., Ke, X., Jeong, B.-C., Wang, Z., Yoo, K.S., Yadav, G.P., Kinch, L.N., et al. (2016a). HP1BP3, a Chromatin Retention Factor for Co-transcriptional MicroRNA Processing. Mol. Cell 63, 420–432.

Liu, Y., Beyer, A., and Aebersold, R. (2016b). On the Dependency of Cellular Protein Levels on mRNA Abundance. Cell 165, 535–550.

Makeyev, E.V., Zhang, J., Carrasco, M.A., and Maniatis, T. (2007). The MicroRNA miR-124 Promotes Neuronal Differentiation by Triggering Brain-Specific Alternative Pre-mRNA Splicing. Mol. Cell 27, 435–448.

Mauger, O., Lemoine, F., and Scheiffele, P. (2016). Targeted Intron Retention and Excision for Rapid Gene Regulation in Response to Neuronal Activity. Neuron 92, 1266–1278.

Naftelberg, S., Schor, I.E., Ast, G., and Kornblihtt, A.R. (2015). Regulation of alternative splicing through coupling with transcription and chromatin structure. Annu. Rev. Biochem. 84, 165–198.

Naganuma, T., Nakagawa, S., Tanigawa, A., Sasaki, Y.F., Goshima, N., and Hirose, T. (2012). Alternative 3′-end processing of long noncoding RNA initiates construction of nuclear paraspeckles. EMBO J. 31, 4020–4034.

Ninomiya, K., Kataoka, N., and Hagiwara, M. (2011). Stress-responsive maturation of Clk1/4 pre-mRNAs promotes phosphorylation of SR splicing factor. J. Cell Biol. 195, 27–40.

Novikov, A.V. (2019). PyClustering: Data Mining Library. J. Open Source Softw. 4, 1230.

Pandya-Jones, A., and Black, D.L. (2009). Co-transcriptional splicing of constitutive and alternative exons. RNA 15, 1896–1908.

Pandya-Jones, A., Bhatt, D.M., Lin, C.-H., Tong, A.-J., Smale, S.T., and Black, D.L. (2013). Splicing kinetics and transcript release from the chromatin compartment limit the rate of Lipid A-induced gene expression. RNA 19, 811–827.

Pandya-Jones, A., Markaki, Y., Serizay, J., Chitiashvili, T., Mancia Leon, W.R., Damianov, A., Chronis, C., Papp, B., Chen, C.-K., McKee, R., et al. (2020). A protein assembly mediates Xist localization and gene silencing. Nature 1–7.

Parra, M., Booth, B.W., Weiszmann, R., Yee, B., Yeo, G.W., Brown, J.B., Celniker, S.E., and Conboy, J.G. (2018). An important class of intron retention events in human erythroblasts is regulated by cryptic exons proposed to function as splicing decoys. RNA N. Y. N 24, 1255–1265.

Parra, M.K., Zhang, W., Vu, J., DeWitt, M.A., and Conboy, J.G. (2020). Antisense targeting of decoy exons can reduce intron retention and increase protein expression in human erythroblasts. RNA rna.075028.120.

Pawlicki, J.M., and Steitz, J.A. (2008). Primary microRNA transcript retention at sites of transcription leads to enhanced microRNA production. J. Cell Biol. 182, 61–76.

Pelleg, D., and Moore, A. (2000). X-means: Extending K-means with Efficient Estimation of the Number of Clusters. In In Proceedings of the 17th International Conf. on Machine Learning, (Morgan Kaufmann), pp. 727–734.

Popp, M.W.-L., and Maquat, L.E. (2013). Organizing Principles of Mammalian Nonsense-Mediated mRNA Decay. Annu. Rev. Genet. 47, 139–165.

Quinn, J.J., and Chang, H.Y. (2016). Unique features of long non-coding RNA biogenesis and function. Nat. Rev. Genet. 17, 47–62.

Sakabe, N.J., and de Souza, S.J. (2007). Sequence features responsible for intron retention in human. BMC Genomics 8, 59.

Saldi, T., Cortazar, M.A., Sheridan, R.M., and Bentley, D.L. (2016). Coupling of RNA Polymerase II Transcription Elongation with Pre-mRNA Splicing. J. Mol. Biol. 428, 2623–2635.

Schmid, M., and Jensen, T.H. (2018). Controlling nuclear RNA levels. Nat. Rev. Genet. 19, 518–529.

Schmitz, U., Pinello, N., Jia, F., Alasmari, S., Ritchie, W., Keightley, M.-C., Shini, S., Lieschke, G.J., Wong, J.J.-L., and Rasko, J.E.J. (2017). Intron retention enhances gene regulatory complexity in vertebrates. Genome Biol. 18, 216.

Seila, A.C., Calabrese, J.M., Levine, S.S., Yeo, G.W., Rahl, P.B., Flynn, R.A., Young, R.A., and Sharp, P.A. (2008). Divergent Transcription from Active Promoters. Science 322, 1849–1851.

Shen, S., Park, J.W., Lu, Z., Lin, L., Henry, M.D., Wu, Y.N., Zhou, Q., and Xing, Y. (2014). rMATS: Robust and flexible detection of differential alternative splicing from replicate RNA-Seq data. Proc. Natl. Acad. Sci. 111, E5593–E5601.

Sibley, C.R., Emmett, W., Blazquez, L., Faro, A., Haberman, N., Briese, M., Trabzuni, D., Ryten, M., Weale, M.E., Hardy, J., et al. (2015). Recursive splicing in long vertebrate genes. Nature 521, 371–375.

Spellman, R., Llorian, M., and Smith, C.W.J. (2007). Crossregulation and Functional Redundancy between the Splicing Regulator PTB and Its Paralogs nPTB and ROD1. Mol. Cell 27, 420–434.

Stewart, M. (2019). Polyadenylation and nuclear export of mRNAs. J. Biol. Chem. 294, 2977–2987.

Tilgner, H., Knowles, D.G., Johnson, R., Davis, C.A., Chakrabortty, S., Djebali, S., Curado, J., Snyder, M., Gingeras, T.R., and Guigó, R. (2012). Deep sequencing of subcellular RNA fractions shows splicing to be predominantly co-transcriptional in the human genome but inefficient for lncRNAs. Genome Res. 22, 1616–1625.

Vargas, D.Y., Shah, K., Batish, M., Levandoski, M., Sinha, S., Marras, S.A.E., Schedl, P., and Tyagi, S. (2011). Single-Molecule Imaging of Transcriptionally Coupled and Uncoupled Splicing. Cell 147, 1054–1065.

Vigot, R., Barbieri, S., Bräuner-Osborne, H., Turecek, R., Shigemoto, R., Zhang, Y.-P., Luján, R., Jacobson, L.H., Biermann, B., Fritschy, J.-M., et al. (2006). Differential Compartmentalization and Distinct Functions of GABAB Receptor Variants. Neuron 50, 589–601.

Vilborg, A., and Steitz, J.A. (2017). Readthrough transcription: How are DoGs made and what do they do? RNA Biol. 14, 632–636.

Vuong, C.K., Black, D.L., and Zheng, S. (2016). The neurogenetics of alternative splicing. Nat. Rev. Neurosci. 17, 265–281.

Wang, Q., and Rio, D.C. (2018). JUM is a computational method for comprehensive annotation-free analysis of alternative pre-mRNA splicing patterns. Proc. Natl. Acad. Sci. U. S. A. 115, E8181–E8190.

Wegener, M., and Müller-McNicoll, M. (2018). Nuclear retention of mRNAs - quality control, gene regulation and human disease. Semin. Cell Dev. Biol. 79, 131–142.

Windhager, L., Bonfert, T., Burger, K., Ruzsics, Z., Krebs, S., Kaufmann, S., Malterer, G., L’Hernault, A., Schilhabel, M., Schreiber, S., et al. (2012). Ultrashort and progressive 4sU-tagging reveals key characteristics of RNA processing at nucleotide resolution. Genome Res. 22, 2031–2042.

Wong, J.J.-L., Ritchie, W., Ebner, O.A., Selbach, M., Wong, J.W.H., Huang, Y., Gao, D., Pinello, N., Gonzalez, M., Baidya, K., et al. (2013). Orchestrated intron retention regulates normal granulocyte differentiation. Cell 154, 583–595.

Wuarin, J., and Schibler, U. (1994). Physical isolation of nascent RNA chains transcribed by RNA polymerase II: evidence for cotranscriptional splicing. Mol. Cell. Biol. 14, 7219–7225.

Yap, K., Lim, Z.Q., Khandelia, P., Friedman, B., and Makeyev, E.V. (2012). Coordinated regulation of neuronal mRNA steady-state levels through developmentally controlled intron retention. Genes Dev. 26, 1209–1223.

Yap, K., Mukhina, S., Zhang, G., Tan, J.S.C., Ong, H.S., and Makeyev, E.V. (2018). A Short Tandem Repeat-Enriched RNA Assembles a Nuclear Compartment to Control Alternative Splicing and Promote Cell Survival. Mol. Cell 72, 525–540.e13.

Yeom, K.-H., and Damianov, A. (2017). Methods for Extraction of RNA, Proteins, or Protein Complexes from Subcellular Compartments of Eukaryotic Cells. In MRNA Processing: Methods and Protocols, Y. Shi, ed. (New York, NY: Springer), pp. 155–167.

Yeom, K.-H., Mitchell, S., Linares, A.J., Zheng, S., Lin, C.-H., Wang, X.-J., Hoffmann, A., and Black, D.L. (2018). Polypyrimidine tract-binding protein blocks miRNA-124 biogenesis to enforce its neuronal-specific expression in the mouse. Proc. Natl. Acad. Sci. 115, E11061–E11070.

Zheng, S., Gray, E.E., Chawla, G., Porse, B.T., O’Dell, T.J., and Black, D.L. (2012). PSD-95 is post-transcriptionally repressed during early neural development by PTBP1 and PTBP2. Nat. Neurosci. 15, 381–388.

